# Distinct algorithms for combining landmarks and path integration in medial entorhinal, visual and retrosplenial cortex

**DOI:** 10.1101/2020.10.05.327106

**Authors:** Malcolm G. Campbell, Alexander Attinger, Samuel A. Ocko, Surya Ganguli, Lisa M. Giocomo

## Abstract

During navigation, animals estimate their position using path integration and landmarks, engaging many brain areas. Whether these areas follow specialized or universal cue integration principles remains unknown. Here, we combined electrophysiology with virtual reality to quantify cue integration across thousands of neurons in three areas that support navigation: primary visual (V1), retrosplenial (RSC) and medial entorhinal cortex (MEC). Path integration influenced position estimates in MEC more than in V1 and RSC. V1 coded position retrospectively, likely reflecting delays in sensory processing, whereas MEC coded position prospectively, and RSC was intermediate between the two. In combining path integration with landmarks, MEC showed signatures of Kalman filtering, and we report a distance-tuned neural population that could implement such filtering through attractor dynamics. Our results show that during navigation, MEC serves as a specialized cortical hub for reconciling path integration and landmarks to estimate position and suggest an algorithm for calculating these estimates.

## Introduction

In mammals, navigation engages multiple brain regions, ranging from primary sensory areas to higher order associative areas. This range supports the representation and integration of the multiple cues encountered during navigation that allow animals to generate an internal estimate of their position in external space. Each spatial location corresponds to a set of external sensory cues, which together can serve as landmarks that inform the animal about its position. The animal can also estimate its position by tracking self-motion over time, even in the absence of landmarks. As they navigate through the world, animals must appropriately integrate these two sources of information – external input from landmark cues and internally generated path integration predictions based on self-motion cues – to accurately compute their position. However, while recent work points to strong commonalities in the representation of sensory and behavioral variables across multiple cortical areas (Allen et al., 2019; Clancy et al., 2019; Minderer et al., 2019; Musall et al., 2019; Pinto et al., 2019; Stringer et al., 2019), the degree to which cortical areas involved in navigation follow common or specialized algorithms for computing position remains incompletely understood.

To address this question, we considered how the representation of an animal’s position is computed in three interconnected brain regions hypothesized to provide complementary computations for visually-guided navigation in mice: the medial entorhinal cortex (MEC), primary visual cortex (V1), and retrosplenial cortex (RSC) (Miller and Vogt, 1984; Sugar et al., 2011). Common across these three regions are neural representations for an animal’s external spatial position during visually-guided navigation and strong modulation of neural activity by locomotion (Alexander and Nitz, 2015; Clancy et al., 2019; Diehl et al., 2017; Fischer et al., 2020; Fiser et al., 2016; Fournier et al., 2020; Hafting et al., 2005; Hardcastle et al., 2017; Keller et al., 2012; Kropff et al., 2015; Mao et al., 2017; Niell and Stryker, 2010; Saleem et al., 2013; Saleem et al., 2018). Moreover, during visually guided navigation, MEC, V1 and RSC have all been proposed to implement algorithms for reconciling actual sensory input with internal models of predicted sensory feedback or spatial position (Alexander and Nitz, 2015; Campbell et al., 2018; Keller et al., 2012). However, MEC, V1 and RSC also show specialization in their functional coding properties observed in behaving animals. MEC neurons encode an animal’s spatial position and orientation in a world-centered (i.e. allocentric) reference frame (Diehl et al., 2017; Hafting et al., 2005; Høydal et al., 2019; Sargolini et al., 2006; Solstad et al., 2008). V1 neurons respond to specific visual stimuli and predicted visual feedback (Hubel and Wiesel, 1962; Keller et al., 2012). RSC neurons conjunctively represent the animal’s spatial position in both world-centered (i.e. allocentric) and self-centered (i.e. egocentric) reference frames (Alexander and Nitz, 2015, 2017; Mao et al., 2017). Thus, a key unknown is the degree to which algorithms for cue combination in each region follow specialized versus shared principles.

One challenge in addressing the degree to which cue combination principles differ across cortical regions has been the diversity of navigational tasks used to examine neural activity in MEC, V1 and RSC. Moreover, recent work has revealed heterogeneity and state-dependent modulation of neural activity in all three of these brain regions (Alexander and Nitz, 2015; Clancy et al., 2019; Hardcastle et al., 2017; Niell and Stryker, 2010), necessitating large-scale simultaneous recordings of many neurons, to account for individual trial, session and animal variability. Here, we addressed these challenges by using Neuropixels probes to simultaneously record from tens to hundreds of MEC, V1 or RSC neurons as mice navigated virtual reality environments in which we systematically perturbed the visual cues. Specifically, we introduce alterations in the relationship between vision and locomotion to dissect the differential contribution of each information source to internal estimates of position across all three brain regions. We found that path integration influenced MEC more than RSC and V1, where position estimates were primarily driven by visual landmarks. V1 and RSC represented position tens of milliseconds in the past, possibly due to delays in sensory processing, whereas MEC represented position tens of milliseconds into the future, using path integration calculations to construct short time scale predictions. More than V1 or RSC, MEC resolved small conflicts between landmarks and path integration following the principles of Kalman filtering. These unique features of MEC were strongest in putative MEC grid cells, identified using recordings in the dark in which they fired periodically with respect to distance, which could reflect the computation of approximate Kalman filtering through attractor network dynamics. Together, these results point to MEC as a specialized circuit for integrating multiple noisy sources of position information during navigation.

## Results

### Differential impact of visual cues on spiking activity in MEC, V1, and RSC during navigation

To explore how the brain combines visual landmarks and path integration during navigation, we trained head-fixed mice to traverse a virtual reality (VR) linear track for water rewards (Figure 1A,B). The track contained 5 visual landmarks (towers) spaced at 80 cm intervals. Mice received a water reward (~2μL) at the tower at 400 cm, at which point they instantly teleported to the beginning of the track for the next trial, giving the impression of running along an infinite hallway. After training (~2 weeks, >1,000 trials) mice developed stereotyped behavior in which they slowed down and licked prior to the reward tower (Figure 1C). Mice then performed this task after acute implantation of a silicon probe, which allowed us to record from thousands of single cells (i.e. units) from MEC, V1 and RSC (MEC cell n = 14,799, session n= 90, mouse n = 20; V1 cell n = 1,524, session n = 24, mouse n = 9; RSC cell n = 1,912, session n = 18, mouse n = 9) (Figure 1D,E, Figure S1A). The cell yield per contact did not differ between brain regions (one-way ANOVA, F(2,133)=2.5, p>0.085) but due to the correspondence between the geometry of the probe and MEC anatomy, the number of units per recording session was highest for MEC (Mean ± SEM; MEC: 166 ± 11 cells/session, V1: 65 ± 7 cells/session, RSC: 73 ± 13 cells/session, one-way ANOVA, F(2,133)=18.79, p<6.7e-8, MEC different from V1: p<6.9*10e-7, MEC different from RSC: p<5.8e-6, Mann-Whitney U test) (Figure 1F). Consistent with a previous report (Hernández-Pérez et al., 2020), population activity traveled from dorsal to ventral MEC in theta-paced waves, whereas we did not observe such activity in RSC or V1 (Figure S1B).

**Figure 1:**
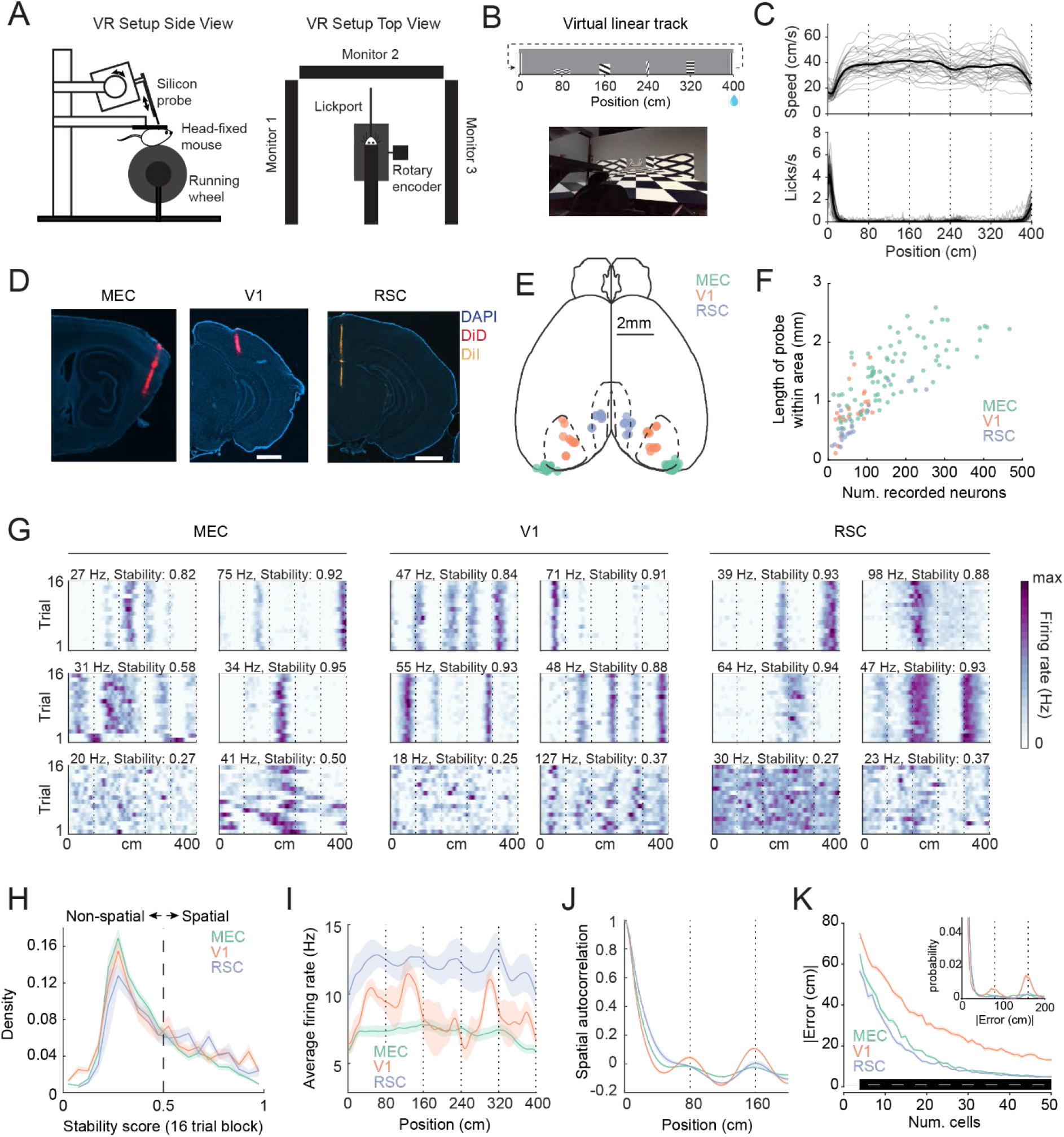
Differential impact of visual landmarks on spiking activity in MEC, V1, and RSC. **A)** Virtual reality setup. Mice were head-fixed and placed on a running wheel. Visual stimuli were presented on 3 monitors arranged in a U-shape around the mouse. The micro-manipulator was placed to maximize unimpeded field of view. Water rewards were delivered via a lick spout. **B)** Top: Schematic of virtual linear track. The track featured five distinct visual landmarks (towers) spaced 80 cm apart. Towers were identical on the left and right and were distinguishable based on width, height and pattern. Mice received a water reward at 400 cm and then teleported to the beginning of the track for the next trial. Bottom: View from behind a mouse. **C)** Average running speed (top) and lick rate (bottom) for all 32 mice. Dotted lines indicate tower locations. Mice slowed down and started licking when approaching the reward tower, indicating they had learned the task. **D)** Example histology images showing probe tracks in MEC, V1, and RSC. The locations of recorded cells (units) were inferred based on histological reconstructions of probe tracks. **E)** Schematic of dorsal brain surface with location of probe insertion on the cortical surface for all 137 recording sessions in 32 mice. Color of dot indicates the targeted brain region. Dashed lines indicate borders of V1 and RSC. **F)** Number of well-isolated units in each recording session. Only units within the target brain area were included. The number of recorded units depended on the length of probe within the targeted area, as recording density was similar across areas. **G)** Spatial firing rate maps across blocks of 16 baseline trials for three example units each in MEC, V1, and RSC. Dashed lines indicate location of towers. The stability score of each neuron (see Methods) for the given block of trials is listed above the rate map. **H)** Average histogram of stability scores across blocks of 16 baseline trials. Stability scores were higher in V1 and RSC than MEC (one-way ANOVA, F(2,96)=5.15, p<0.0075, Stab_MEC_<Stab_V1_: p<0.016, StabMEC<StabRSC: p<0.002, one-sided Mann-Whitney U test). Cells were defined as ‘spatial’ within a given trial block if they had stability score ≥ 0.5. Values were averaged within session first, then across sessions (error bars are over sessions, not cells). **I)** Average firing rate of all spatial cells in MEC (green), V1 (red), and RSC (blue) with respect to track location. Dashed lines show tower locations. V1 and RSC activity was modulated by towers, whereas MEC activity was relatively uniform with respect to the track. RSC firing rates were significantly higher overall (one-way ANOVA, F(2,94)=12.17, FRRSC>FRV1: p<1.4*10e-4, FRRSC>FRMEC: p<0.005, one-sided Mann-Whitney U test). Values were averaged within session first, then across sessions. **J)** Average spatial autocorrelation for all spatial cells. V1 and RSC, and to a lesser extent MEC, show peaks at lags corresponding to distance between adjacent towers. Values were averaged within session first, then across sessions. **K)** Position decoding error as a function of the number of included spatial neurons (Methods). Inset shows the distribution of errors for a decoder trained on 20 neurons. V1 errors were larger than MEC and RSC errors because of frequent tower misclassifications in which the difference between the true and decoded position differed by a multiple of 80 cm (inset). Black bar at bottom indicates where p<0.05 for ErrorV1>ErrorMEC and ErrorV1>ErrorRSC, Mann-Whitney U test.

In MEC, V1 and RSC, many cells fired action potentials at consistent positions along the VR track. We defined such cells as ‘spatially stable’ and identified them using a cross correlation metric (Methods). Averaging the firing rates of all spatially stable cells across the VR track revealed clear population level firing pattern differences between the three brain regions. In V1 and RSC, firing rates peaked ~20 cm before each visual landmark (Figure 1H-J). In MEC, firing rates uniformly tiled the entire VR track (Figure 1H-J). Even so, the position of the mouse on the VR track could be decoded with high accuracy from the neural activity of any of the three brain regions (Figure 1K). However, we observed more error in decoded position estimates from V1 neural activity. This decoded error emerged from misclassified tower locations, possibly due to similar population level firing patterns in V1 near visual landmarks, leading to the increased prevalence of 80 cm and 160 cm errors (Figure 1K, inset, Figure S1C). Together, these analyses reveal that neural representations in MEC, V1 and RSC all carry information regarding an animal’s spatial position but that visual landmark cues influence these neural representations in V1 and RSC more than MEC. Nevertheless, visual landmarks were necessary, but not sufficient, to drive spatial firing patterns in MEC (Figure S1D-G).

### Conflicts between landmarks and path integration change neural activity patterns in MEC more than in V1 or RSC

We next examined the degree to which visual landmarks versus path integration determined neural activity patterns in MEC, V1 and RSC. We created visuomotor cue conflicts by reducing the gain between the rotation of the running wheel and the mouse’s progression along the virtual hallway by a factor of 0.8 for blocks of 4 trials (number of blocks per session = 1-4, median 2) (Figure 2A). This gain manipulation had minimal effects on the animals’ behavior (Figure 2B), and no effect on average firing rates (Figure S2A). If the location of spikes is determined solely by the visually determined position of the mouse in the virtual environment, neural activity patterns should not shift positions between baseline and gain trials. Conversely, if position estimates are influenced by actual distance run on the treadmill due to self-motion, then neural activity patterns will undergo negative (leftward) shifts when plotted as a function the visually determined position in the environment, since the visual environment now advances less for a given number of steps taken. Alternatively, gain changes could trigger global remapping, which was previously shown to occur in MEC grid cells for large enough conflicts between path integration and landmarks (Campbell et al., 2018).

**Figure 2:**
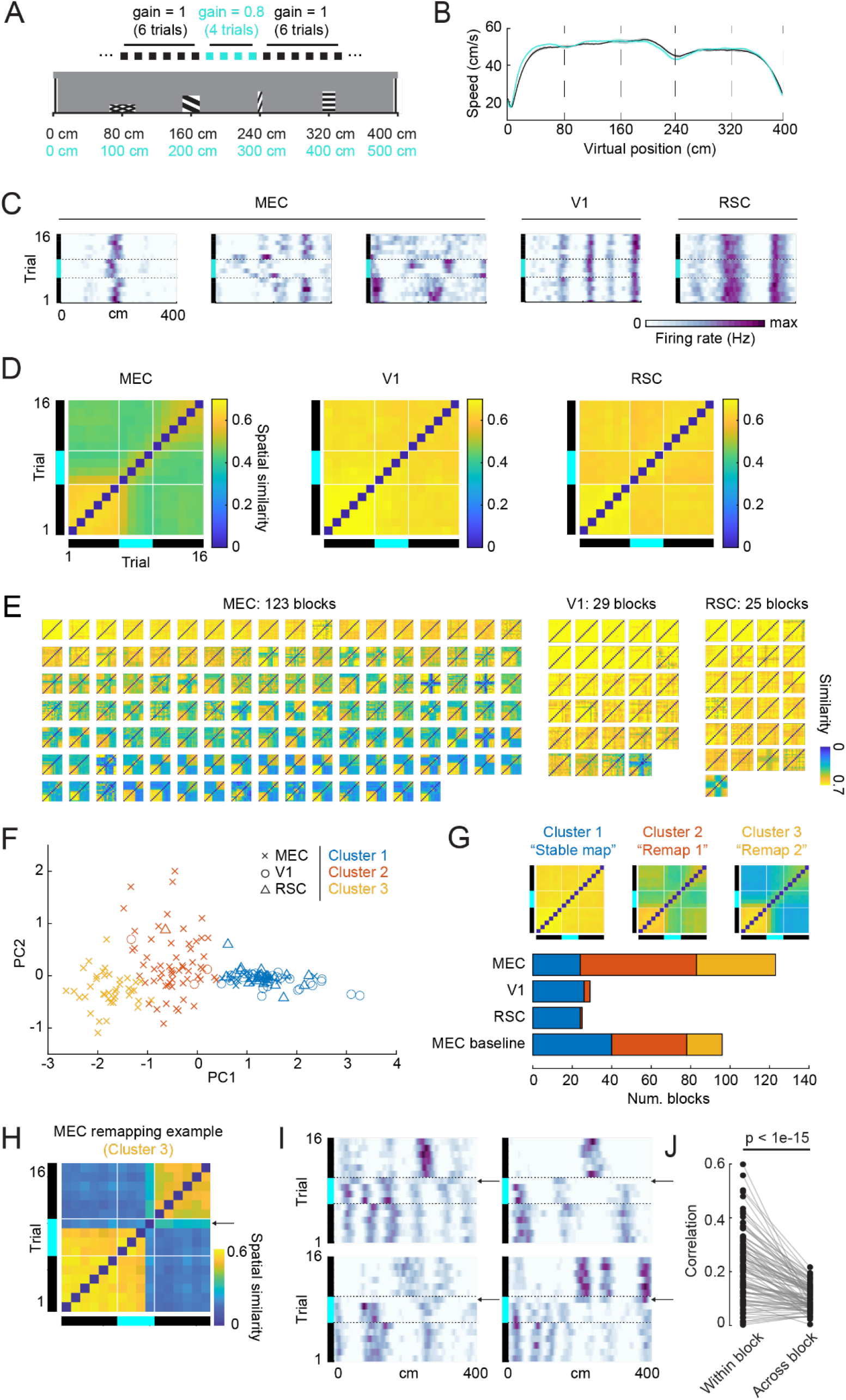
Conflicts between landmarks and path integration change neural activity patterns in MEC more than in V1 or RSC. **A)** Trial structure for gain change experiments. Baseline trials (black) were interleaved with blocks of four gain change trials (cyan). **B)** Average running speed along virtual reality track during baseline (black) and gain change trials (cyan). Dotted lines indicate the position of visual landmarks. Average running speed did not differ between the two conditions (average speed ± SEM: baseline = 47.0 ± 9.7 cm/s, gain change = 47.4 ± 9.2 cm/s, p = 0.167, 216 gain change blocks, Wilcoxon test). **C)** Spatial firing rate maps for 16 trials in and surrounding gain change blocks for example MEC, V1, and RSC neurons. Baseline trials are indicated by black bar, gain change trials are indicated by cyan bar. Whereas firing patterns were mostly unchanged in V1 and RSC during gain manipulations (right), neurons often remapped in MEC (left). **D)** Trial-trial similarity matrices for each brain region, averaged over gain change blocks (see Methods). Black and cyan bars indicate baseline and gain change trials, respectively. White lines separate gain change trials. **E)** Average trial-trial similarity matrices as in (D) but each block represents a trial-trial similarity matrix for an individual gain change block. The average similarity matrices in (D) obscured substantial variability in the degree and pattern of MEC remapping (left), whereas spatial firing patterns of V1 and RSC neurons were more stable and rarely remapped (right). **F)** Projection onto first two principal components for each similarity matrix. Type of marker indicates brain region, color of marker indicates cluster assignment of k-means clustering (see Methods). **G)** Top: Average similarity matrices for the three clusters found in (F). The three clusters roughly corresponded to the three example MEC neurons shown in (C). Bottom: Number of gain change blocks assigned to each cluster by brain region. For comparison, we ran the same analysis on MEC baseline data in which no gain change occurred. Remapping could occur spontaneously in MEC (see also Figure S2J). However, remapping was significantly more frequent in response to a gain change (Fisher test p = 8.9e-4). **H)** Similarity matrix of an example Cluster 3 MEC gain change block. Neurons remap in gain change trial 4 (arrow), leading to low pairwise similarity between trials before and after remapping. **I)** Spatial firing rate maps of four example MEC neurons during the gain change block shown in (H). During the first three gain change trials, (black bar: baseline, blue bar: gain change, dotted line separates gain change / baseline trials), the spatial maps are shifted relative to the preceding baseline map. The neurons remap in gain change trial 4 (arrows). Spatial maps in post-baseline trials (trials 11-16) are different from the spatial maps in baseline trials. **J)** For each MEC trial block in cluster 2 and cluster 3 (99 trial blocks total), we assessed coordination of remapping among simultaneously recorded spatial cells by comparing the correlation of spatial similarity maps for simultaneously recorded spatial cells, to all other similarity maps (see Methods). Similarity maps of simultaneously recorded were more similar to each other than to similarity maps of other groups of units, consistent with population wide remapping (p<=8.2*10e-16, Wilcoxon signed rank test).

We considered spatially stable cells, classified as cells with an average peak trial-trial firing rate correlation ≥ 0.5 in the 6 baseline trials preceding the gain change (Methods). In MEC, cells responded to the change in gain in one of two ways: the spatial firing pattern of a cell shifted coherently, or the firing pattern remapped, consistent with previous work (Campbell et al., 2018). When cells remapped in response to the gain change, they either returned to the original baseline map following the gain change, or did not (Figure 2C). In contrast, firing patterns in V1 and RSC remained remarkably stable both during and after the gain change (Figure 2C), consistent with the idea that visual cues strongly influenced firing patterns in these brain regions (Figure 1I).

To quantify the effect of the gain change on neural firing patterns, we computed similarity matrices by correlating firing patterns on pairs of trials (Figure 2D, Methods). We averaged the similarity matrix over cells for a gain change block and considered each block of trials separately, as remapping patterns and the number of spatially stable cells varied across blocks (MEC: trial block n = 123, session n = 51, mouse n = 18, 51 ± 3 (SEM) cells per block; V1: trial block n = 29, session n = 15, mouse n = 6, 20 ± 2 (SEM) cells per block; RSC: trial block n = 25, session n = 13, mouse n = 7, 55 ± 8 (SEM) cells per block) (Figure 2E; for cell-level results see Figure S2B,C). In MEC, we observed variability in the similarity matrices (Figure 2E, left), whereas in V1 and RSC these matrices almost always indicated that neural representations remained stable (did not remap) between baseline, gain change and post-gain change trials (Figure 2E, right, Figure S2D-F). To identify distinct types of population-level gain change responses, we reduced the dimensionality of these data using PCA and then applied k-means clustering (k=3) (Figure 2F, Figure S2G). The resulting clusters corresponded to three types of responses between baseline, gain change and post-gain change trials: the neural population did not remap across conditions (“Stable”, Cluster 1), the neural population remapped during the gain change but returned to the baseline map in the post-gain change trials (“Remapping 1”, Cluster 2), or the neural population remapped during the gain change and did not return to the baseline map in the post-gain change trials (“Remapping 2”, Cluster 3) (Figure 2G). In MEC, many trial blocks were assigned to Clusters 2 or 3, indicating cue conflict often led to remapping of spatial firing patterns (blocks in cluster/total blocks: MEC, cluster 1 = 24/120, cluster 2 = 59/120, cluster 3 = 40/120). This gain change-induced remapping was remarkably coordinated across neurons across the entire dorsal ventral extent of MEC (Figure 2H-J), with remapping patterns more similar within block than between blocks (Figure 2J). We also observed occasional spontaneous remapping in MEC in the absence of any gain change (blocks in cluster/total baseline blocks: cluster 1 = 40/96, cluster 2 = 38/96, cluster 3 = 18/96) (Figure 2G, Figure S2H-J) (Low et al., BioRXiv 2020). However, the frequency of remapping was significantly higher during gain change blocks than baseline blocks (Fisher test p = 0.00089). In contrast to the variability in MEC responses, in V1 and RSC, nearly all trial blocks were assigned to Cluster 1 (blocks in cluster/total blocks: V1, cluster 1 = 26/29, cluster 2 = 3/29, cluster 3 = 0/29; RSC, cluster 1 = 24/25, cluster 2 = 1/25, cluster 3 = 0/25). This suggests that in V1 and RSC, visual landmarks drive spatial firing patterns, even when visual landmark and path integration cues are in conflict.

### Even in the absence of remapping, path integration influences position estimates in MEC more than in V1 or RSC

While we observed variability in how the MEC neural population responded to the gain change, namely that spatial maps either remained stable or remapped, we next focused on “stable” trial blocks (Cluster 1) in order to compare cue integration mechanisms across brain regions (MEC: 24 Cluster 1 blocks, (mean ± SEM) 65 ± 8 spatial cells per block; V1: 26 Cluster 1 blocks, 21 ± 3 cells per block; RSC: 24 Cluster 1 blocks, 55 ± 9 cells per block; Fig. 3A). In all three brain regions, we observed that even in ‘stable’ blocks, firing rate maps shifted coherently in the direction predicted by path integration on gain change trials; that is, firing rate maps kept their overall pattern but spikes occurred earlier on the VR track (Figure 3B). To quantify the degree of shift between pairs of trials, we computed trial-trial shift matrices by identifying the spatial lag that gave maximum cross-correlation between the trials’ rate maps (Figure 3C). As in Figure 2, we averaged over cells within each trial block and treated each trial block as a single sample. Map shifts on gain change trials were significantly different from zero in all three brain regions (mean map shift ± SEM: MEC = −4.1 ± 0.5 cm, p = 2.1e-5, V1 = −1.8 ± 0.5 cm, p = 0.00022, RSC = −1.4 ± 0.4 cm, p = 0.0047, Wilcoxon tests), but these shifts were largest in MEC (one-way ANOVA, F(2,71)=8.53, p = 0.00048, shifts in MEC larger than V1, p = 0.00057, and RSC, p = 0.00093, Mann-Whitney U test, Figure 3D; for cell-level results see Figure S3A), indicating a stronger influence of path integration on position estimates in MEC compared to V1 or RSC. Moreover, we found that in MEC, but not V1 and RSC, the map shifts of individual neurons systematically deviated from the population average (S3C-E). This indicated that in MEC, cells consistently fell on a spectrum from more landmark-driven to more path integration-driven, whereas in RSC and V1, the relative influence of path integration versus landmarks was constant across cells.

**Figure 3:**
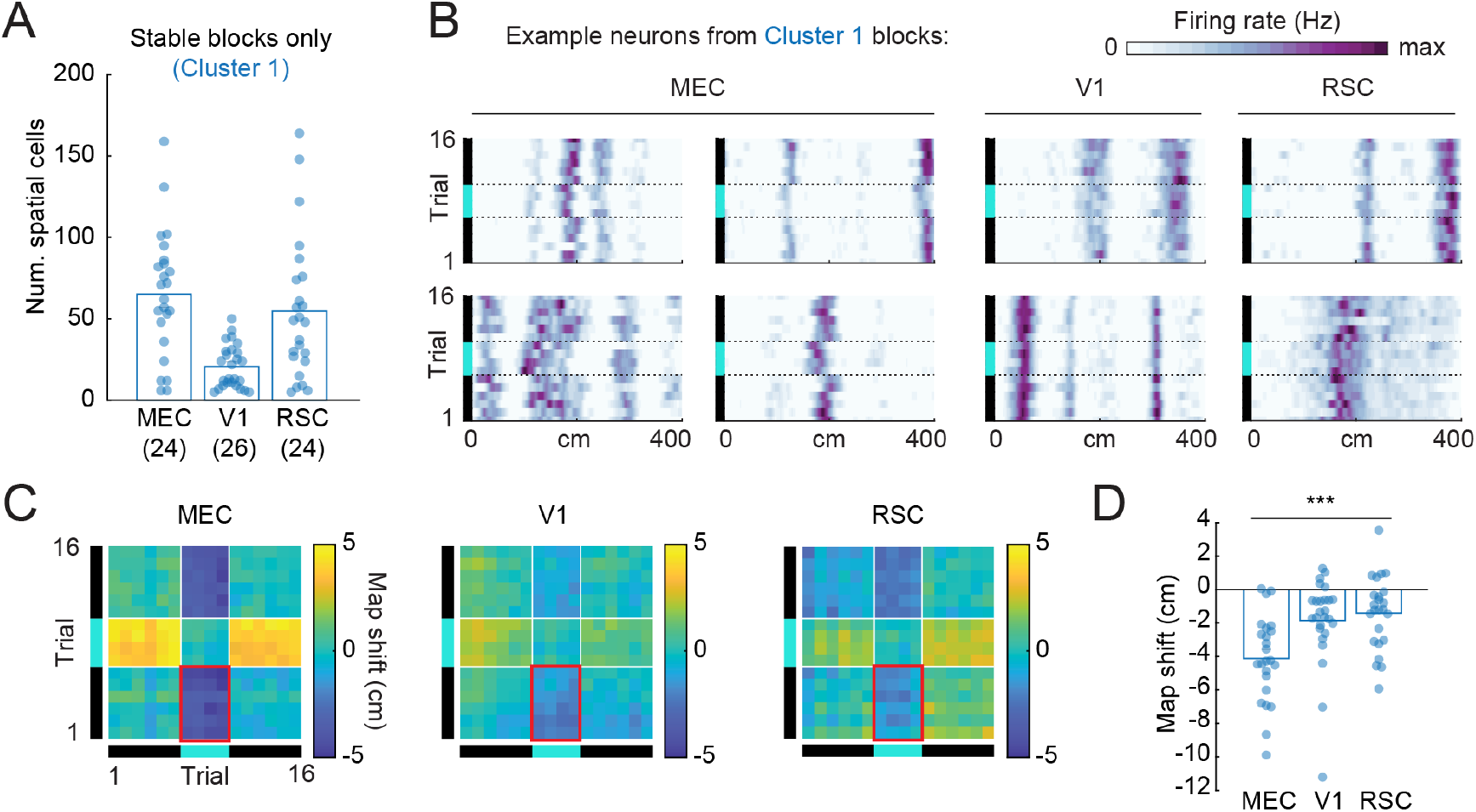
The influence of path integration on position estimates is greater in MEC than in V1 or RSC. **A)** Number of gain change blocks for each brain region identified as “stable” (cluster 1) in Figure 2G. Bars indicate mean. **B)** Example spatial firing rate maps for neurons from cluster 1 blocks in MEC (n = 4), V1 (n = 2), and RSC (n = 2). Note the larger map shifts during gain change trials in MEC compared to V1 and RSC. **C)** Average trial-trial shift matrices for MEC, V1, and RSC. Shifts were computed as the location of the peak spatial cross-correlation for each cell, then averaged over cells within gain change blocks. White lines indicate transition between baseline and gain change trials. Red rectangles indicate samples used for estimating average map shift. **D)** Average map shifts between gain change trials and baseline trials (outlined in red in C) in MEC, V1, and RSC. Each point represents a gain change block. Map shifts were larger in MEC than in V1 or RSC. Bars indicate mean.

### Position is encoded retrospectively in V1 and RSC but prospectively in MEC

Although the gain change-induced map shifts we observed could reflect the influence of path integration, map shifts could also be caused by temporal delays between the encoded position and the time of spikes (Figure 4A). For example, if there exists a delay Δ*t* in sensory processing, a mouse running at velocity *v* when it encounters a visual stimulus will have moved a distance Δ*x* = *v* ⋅ Δ*t* when the first visually driven spikes occur. Such effects could interact with gain changes as a reduction in gain enforces a slower running speed relative to visual cues. We therefore considered the extent to which V1, RSC, and MEC code exact current position versus past or future position (retrospective/prospective coding, respectively) by examining the effect of running speed on spike position in baseline trials. For retrospective coding, spikes will occur later on the track at faster running speeds, and vice versa for prospective coding (Figure 4B). If no retrospective or prospective coding exists, running speed will not influence spike position.

**Figure 4:**
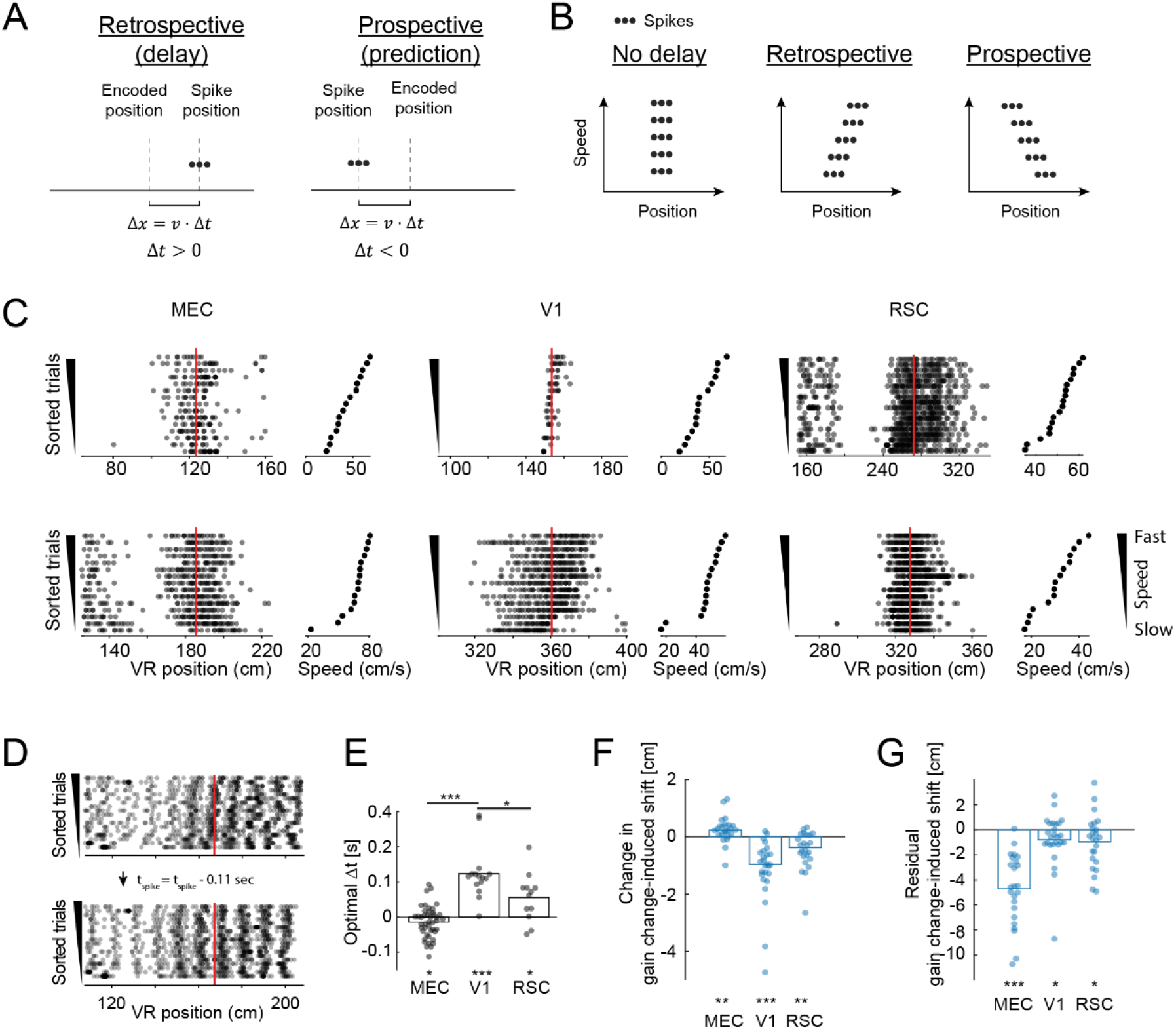
V1 and RSC code position retrospectively whereas MEC codes position prospectively. **A)** Cartoon illustrating retrospective coding (spikes encode past position, left) versus prospective coding (spikes code future position, right). Prospective coding makes use of path integration calculations to predict future position. Retrospective coding can reflect delays in sensory processing. Δ*x*: the spatial offset between the encoded position and the position at which spikes occur. *v*: the mouse’s running speed. Δ*t*: the time delay between the mouse being located at the encoded position and the occurrence of spikes (we define Δ*t* to be negative for prospective coding and positive for retrospective coding). **B)** Schematic showing the predicted influence of running speed on spike position for no delay (left), retrospective (middle), and prospective (right) neural codes. Faster running speeds lead to larger shifts between spike position and encoded position, but in opposite directions for retrospective versus prospective coding. **C)** Example spike raster plots for two MEC, V1, and RSC units for 16 baseline trials sorted by running speed. For each unit, we calculated the location of the maximum average firing rate during baseline trials, and the raster plots are zoomed in to show spikes occurring around the location of the maximum firing rate (indicated by red line). Trials are ordered by the running speed, averaged during 10 cm preceding the location of the peak firing rate. Running speed for each trial is shown on the right of each raster plot. **D)** Illustration showing temporal delay estimation and correction for an example V1 neuron. Top: raster plot with trials ordered by running speed. Note how spikes shift left for slower running speeds. For each unit, we searched for the factor that maximized the trial-by-trial correlation of firing rate maps. For this example unit, estimated temporal delay was +110 ms and we accounted for this delay by changing the time of each spike (equation in middle). After applying this optimal correction, the spikes were more aligned (bottom). **E)** Average estimated temporal delay for each recording, split by brain region. Each dot represents the estimated temporal delay, averaged across all units across a session (MEC: 56 sessions, V1: 18 sessions, RSC: 14 sessions). Estimated temporal delay differed from 0 in all regions (MEC, p<=0.0068, V1: p<=4.2e-4, RSC p<=0.016, Wilcoxon signed rank test). The estimated delay factors were significantly larger in V1 compared to RSC or MEC (one-way ANOVA, F(2,82)=13.25, p<1.3e-5, Delay_V1_ > Delay_MEC_, p<=1.3e-7, Delay_V1_> Delay_RSC_, p<=3.1e-3, Mann-Whitney U test). Bars indicate mean. **F)** Change in map shift after spike times were corrected by the optimal temporal delay factor for each cell. Map shifts increased in MEC (p<=0.0027, Wilcoxon signed rank test), and decreased in V1 and RSC (V1: p<=1.7*10e-5, RSC: p<=0.0015, Wilcoxon signed rank test). **G)** Residual map shifts after correcting for temporal delay, isolating the influence of path integration on position estimates. Residual map shifts in V1 and RSC were non-zero (V1, p<=0.036; RSC, p<=0.032, Wilcoxon signed rank tests).

We considered blocks of baseline trials (no gain change) and sorted these trials by the running speed of the mouse around the location of peak firing rate for each cell separately. In V1 and RSC, the VR location at which spikes occurred often shifted positively with running speed (Figure 4C), consistent with retrospective coding. In contrast, spikes of MEC neurons often shifted in the opposite direction, consistent with prospective coding (Figure 4C). To quantify the shift for each cell, we computed a temporal delay factor for each neuron by maximizing the average trial-trial similarity, which resulted in better alignment of spikes to spatial position across trials (Figure 4D) (Methods). We then averaged across co-recorded neurons to obtain a temporal delay factor for each session. In V1 and, to a lesser extent, RSC, the average delay factor was significantly greater than zero (retrospective coding; mean delay ± SEM: V1 = 0.12 ± 0.02 sec, p < 4.2e-4, RSC = 0.06 ± 0.02 sec, p = 0.016; Figure 4E). In contrast, the average delay factor was negative in MEC, indicating prospective coding (mean delay ± SEM, MEC = −0.01 ± 0.006 sec, p = 0.0068; Figure 4E). A decoding analysis gave qualitatively similar results (Figure S3F).

Consistent with presence of retrospective coding in V1/RSC and prospective coding in MEC, correcting the spike times of individual cells by their delay factor significantly reduced map shifts in V1 and RSC, but increased map shifts in MEC (Figure 4F,G). This supports the idea that, during gain change, temporal delays do not drive the map shifts observed in MEC but do contribute to map shifts in V1 and, to a lesser extent, RSC. Moreover, the prospective coding, combined with large map shifts observed on gain change trials in MEC raises the possibility that MEC uses path integration to predict current position before the arrival of delayed sensory input.

### A simple Kalman filter model for combining path integration and landmarks

We have demonstrated that path integration and landmarks both strongly influence MEC codes for position. How might these sources of information be combined? One algorithm for combining the predictions of an internal model (i.e. path integration estimates) with external inputs (i.e. landmark cues) to optimally estimate an unknown variable (i.e. spatial position) is a Kalman filter (Deneve et al., 2007; Kalman, 1960). To examine the extent to which our neural data are consistent with the algorithmic principle of Kalman filtering, as well as derive further predictions, we implemented a Kalman filter model that combines path integration with external landmark cues (Figure 5A) (Methods).

**Figure 5:**
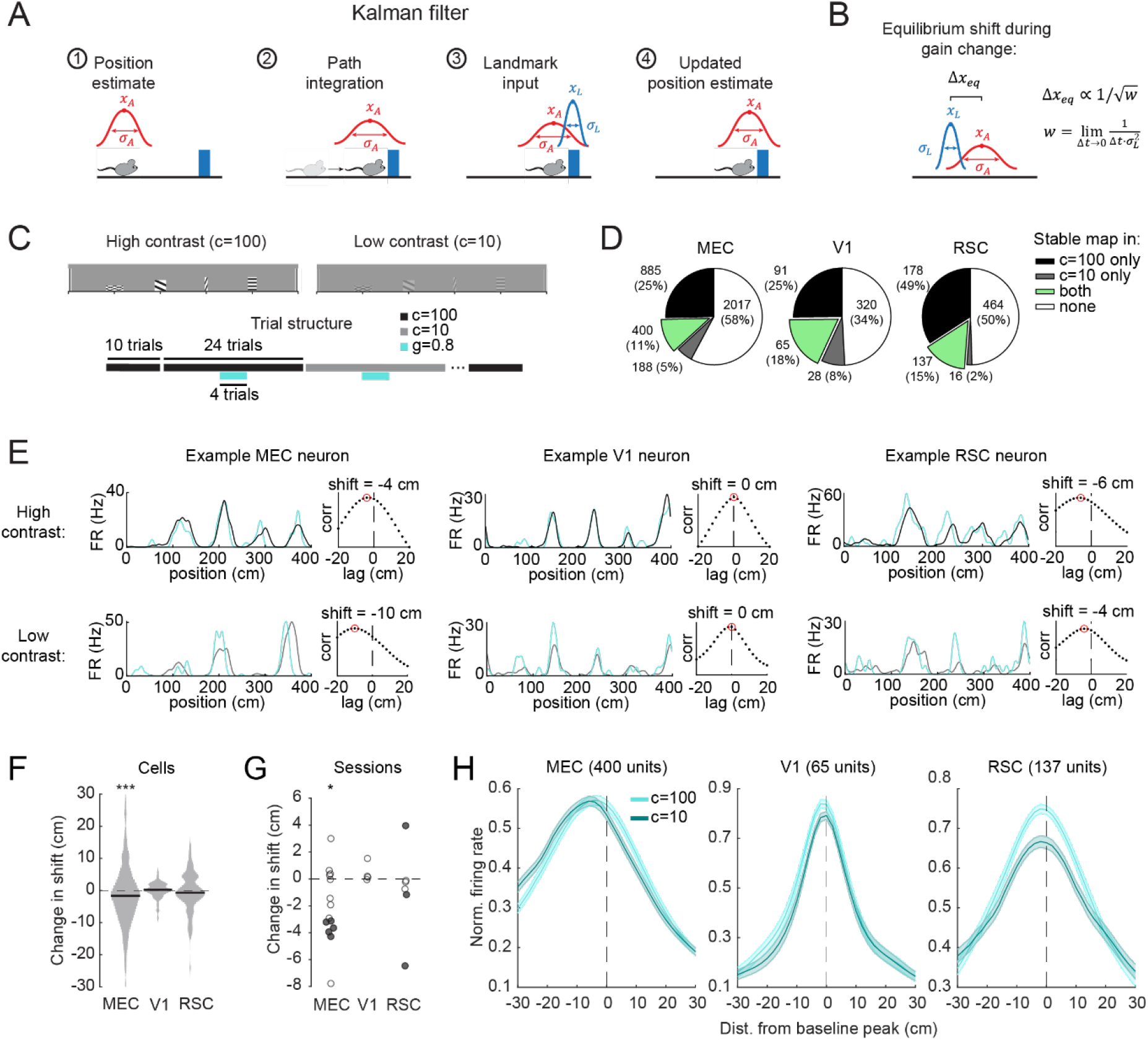
The dependence of map shifts on visual contrast is consistent with a Kalman filter model in MEC but not V1 or RSC. **A)** Cartoon of Kalman filter model that combines path integration and landmarks (see Equations 1–5). *x_A_* is the mouse’s internal estimate of its position, *σ_A_* is the uncertainty of the internal estimate, *x_L_* is the position estimate given by external landmark input, and *σ_L_* is the uncertainty of the landmark input. **B)** During a gain change, the Kalman filter reaches an equilibrium map shift (Δ*x_eq_*) between the internal estimate (*x_A_*) and the landmark input (*x_L_*) which is inversely proportional to the square root of the Kalman landmark strength (*α*) (Methods, Figure S4). The Kalman landmark strength is defined as 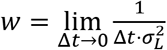 **C)** Experimental design to test the influence of visual contrast (a proxy for *w*) on gain change responses. **D)** Percent of cells with stable gain change responses in high contrast (black), low contrast (grey) or both (green). Roughly 10-20% of cells were stable in both contrast conditions (MEC, 400/3490, 11%; V1, 65/362, 18%; RSC, 137/937, 15%). Only these cells were included in the following analyses, which allowed comparisons between the same cells across contrasts. **E)** Example neuron firing rates in MEC (left), V1 (middle), and RSC (right) during gain changes in high contrast (c=100, top row) and low contrast (c=10, bottom row). Left panels: Black/gray: Average rate map during baseline trials. Cyan: Firing rate map during a single example gain change trial that did not exhibit remapping (peak cross-correlation relative to baseline map ≥ 0.5). Right panels: Cross-correlation between gain change and baseline maps, with the peak indicated by a red circle. The amount of shift was determined by the location of this cross-correlation peak. For each cell, shifts were averaged across trials. **F)** Change in gain change-induced map shifts between high and low contrast for all cells identified in (B). Negative values indicate increased leftward shifts in low contrast. Of the three brain regions, only MEC showed increased shifts in low contrast (change in map shift, c=10-c100, mean ± SEM: MEC = −1.5 ± 0.5 cm, p = 2.7e-5; V1 = 0.0 ± 0.3 cm, p = 0.52; RSC = −0.5 ± 0.5 cm, p = 0.39; Wilcoxon signed rank tests). As a population, shift changes were larger for MEC than V1 and RSC (MEC vs. V1, p = 0.010; MEC vs. RSC, p = 0.037; V1 vs. RSC, p = 0.35; Mann-Whitney U tests). P values were not corrected for multiple comparisons. **G)** To test whether results in (F) were consistent across sessions, we took the median shift change within each session that had at least 5 spatially stable cells in both high and low contrast. The resulting pattern of shift changes was consistent with the cell-level analysis shown in (F), with MEC, but not V1 or RSC, showing significantly larger (more negative) map shifts in low contrast (mean shift change ± SEM over sessions, MEC = −2.0 ± 0.7, N = 14 sessions, p = 0.025; V1 = 0.4 ± 0.4 cm, N = 4 sessions, p = 0.25; RSC = −0.8 ± 1.4 cm, N = 6 sessions, p = 0.31; Wilcoxon signed rank tests). Filled circles indicate sessions with significant within-session shift changes (p < 0.05, Wilcoxon signed rank tests). In MEC, 5/14 sessions showed significant within-session shift changes (larger leftward shift). In V1, 0/4 sessions had significant changes. In RSC, however, there was some diversity amongst sessions: Two sessions showed significantly larger leftward shifts (negative values), whereas another showed significantly smaller leftward shifts (positive values), and three showed no change. **H)** Average normalized firing rate during gain change trials, aligned to the location of peak firing rate in the baseline map. Firing rates were normalized within each trial. Different shades of cyan correspond to the two contrast (c) values. This complementary analysis supports the results of panels (F) and (G), with larger leftward map shifts in low contrast in MEC but not V1 or RSC. *** p < 0.001, ** p < 0.01, * p < 0.05.

The Kalman filter maintains a time-evolving estimate of the animal’s position (*x_A_*) and its uncertainty (*σ_A_*). For simplicity, we only considered mean position estimates and their uncertainties. This allowed us to derive the predicted average neural response to gain manipulations without modeling the random errors that occur on individual trials.

Specifically, at each time step, *x_A_* and *σ_A_* are updated by path integration as follows:

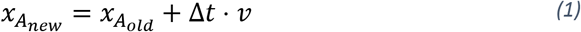

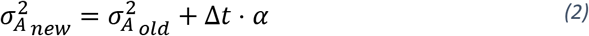

Where Δ*t* is the length of the time step and *α* is a diffusivity constant that describes the rate at which position uncertainty grows during path integration (Methods). Then, within the same time step, landmark input tells the animal it is located at *x_L_* with uncertainty *σ_L_*. The position estimate and its uncertainty are updated using the Kalman gain 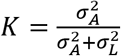:

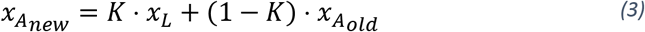

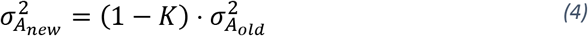

We made the model continuous by taking the limit Δ*t* → 0 (Methods). During a gain manipulation, when the internal velocity estimate (*v_A_*) systematically differs from the landmark velocity (*v_L_*), the Kalman filter reaches an equilibrium shift between the internal position estimate (*x_A_*) and the landmark input (*x_L_*) (Figure 5B):

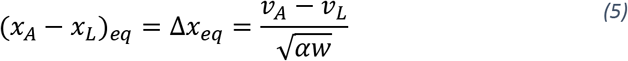

Where 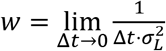 (Methods). We call *w* the “Kalman landmark strength” as it captures the strength (or certainty) of the landmark inputs in the continuous limit.

The shift in the Kalman filter output during gain manipulations resembles the map shifts we observed in MEC in ‘stable’ (cluster 1) blocks (Figure 3). Interestingly, an attractor network model of grid cells can produce similar shifts (Campbell et al., 2018; Ocko et al., 2018) (Figure S4A). Such attractor models capture both the ‘stable’ shifts and ‘remapping’ of MEC neural representations during gain changes (Figure 2, Figure S4B). For small gain changes (*v_A_* ≈ *v_L_*), the network does not remap, and instead reaches an equilibrium shift between the attractor state (*θ_A_*) and the landmark input (*θ_L_*):

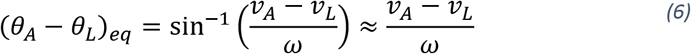

This equation is identical to Equation 5, with *ω* standing in for 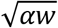. Thus, for small mismatches between path integration and landmark cues, the attractor model is to first approximation equivalent to a Kalman filter. This suggests that attractor dynamics could be used to implement a Kalman filter in MEC.

Note that the Kalman filter as currently described shows how velocity and landmark information can be fused to construct an estimate of current position. To incorporate prospective coding, the model can be easily extended to a Kalman predictor which instead encodes at any given time an extrapolated future position based on current position and velocity. Based on the prospective coding we observe in MEC, we hypothesize MEC instantiates a short time scale Kalman predictor rather than a Kalman filter.

### In MEC but not V1 or RSC, gain change-induced map shifts increase at low visual contrast, consistent with Kalman filtering

We next examined whether neural dynamics in MEC, V1 or RSC were consistent with Kalman filtering. Equation 5 predicts that map shifts observed during gain manipulations depend on the certainty of landmark cues. To test this experimentally, we lowered the contrast of visual cues to reduce the strength (or certainty) of landmark input while also performing gain changes (Figure 5C). If MEC implements Kalman filtering, the decrease in landmark input (a reduction of *w* in Equation 5) increase map shifts. We chose contrast = 10%, as it reduced but did not eliminate MEC map stability (Figure S5). We identified cells with stable maps in both high and low contrast baseline trials (mean baseline peak correlation ≥ 0.5) and selected gain change trials that were sufficiently similar to baseline to exclude remapping (peak correlation to baseline ≥ 0.5; Figure 5D). Consistent with the Kalman filtering hypothesis, lowering the contrast increased map shifts in MEC. On the other hand, map shifts in V1 and RSC did not change (Figure 5E-H). Lowering the contrast to 20% did not influence MEC map shifts (Figure S5H). These results suggest that MEC combines path integration and landmarks following Kalman filtering principles, whereas V1 and RSC are dominated by visual input during the g=0.8 condition.

### A population of putative MEC grid cells tuned to distance run in the dark drives many of MEC’s unique cue integration properties

Given the strong influence of path integration on spatial maps in MEC relative to V1 and RSC, we next aimed to examine if MEC contained pure path integration signals by allowing mice to run on the treadmill in near-total darkness for ~30 minutes (Figure 6A). Many MEC neurons showed striking periodic modulation of their firing rate as a function of distance run in the dark (Figure 6B). The degree of distance tuning exhibited by a cell was determined by computing the autocorrelation of each cell’s firing rate trace, binned by distance run (Figure 6C). Distance tuned cells were then defined as cells with both a maximum autocorrelation peak greater than the 99^th^ percentile of a shuffled distribution and an autocorrelation peak with a prominence value > 0.1 (Figure 6D, Methods). Using these criteria, 11% of all MEC cells were distance-tuned (1603/14172), but percentages within individual sessions ranged from 0-52% (maximum of 111 co-recorded distance cells; Figure 6E, F). Plotting firing rate correlations for all distance cells in individual sessions revealed structured distance tuning that was strikingly reminiscent of MEC grid cells, as preferred distances increased from dorsal to ventral MEC in discrete jumps of roughly a factor of 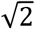 (Figure 6E,G,H,I) (Stensola et al., 2012). Thus, we considered these neurons to be putative MEC grid cells.

**Figure 6:**
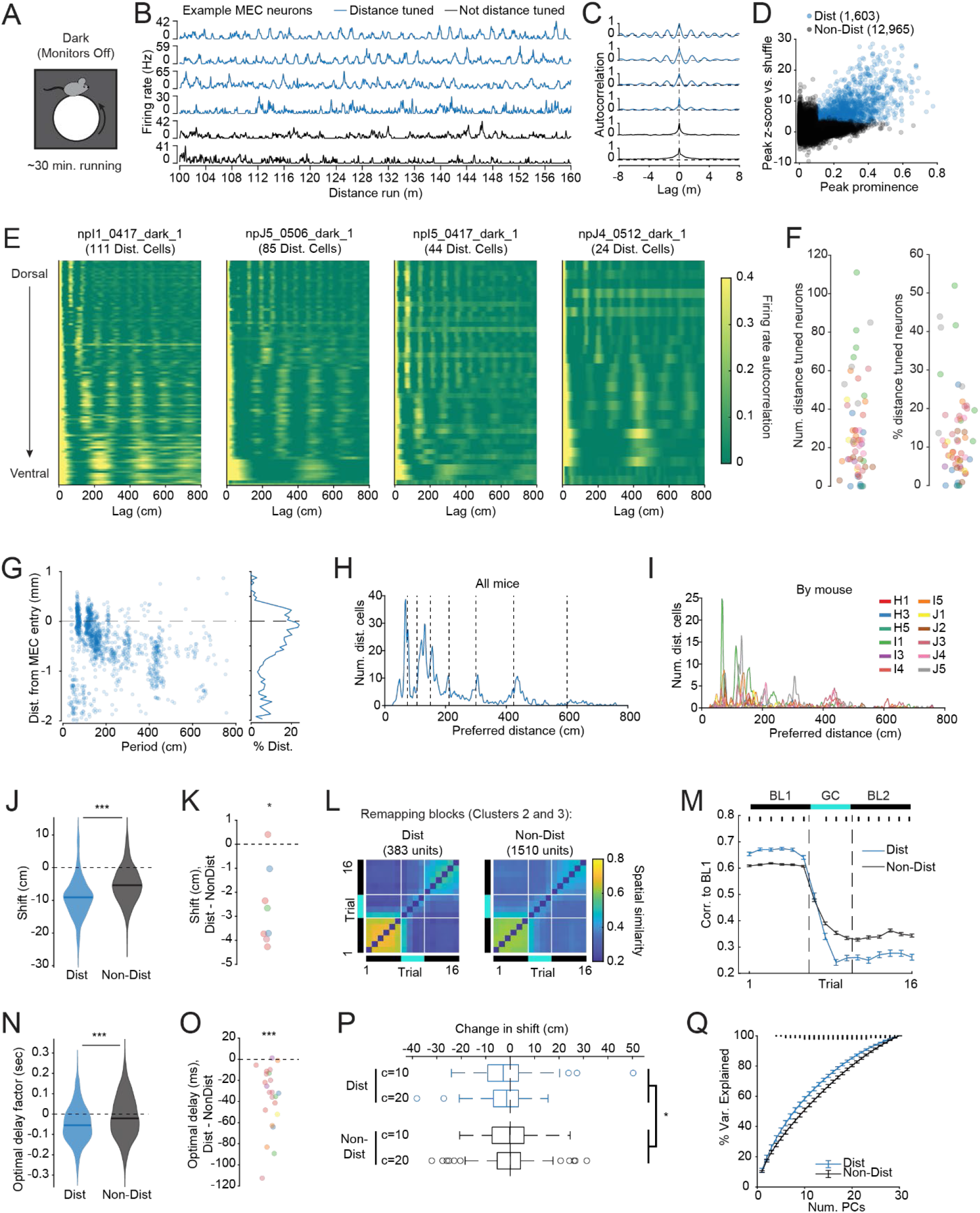
A population of putative MEC grid cells tuned to distance run in the dark drives many of MEC’s unique cue integration properties. **A)** In dark experiments, mice ran freely in near-complete darkness with no rewards for ~30 mins while we recorded MEC neural activity. Dark experiments were typically run at the end of the recording session after we ran one or more VR experiments (Methods). **B)** Example snippet of the firing rate of 6 MEC neurons versus distance run in darkness. A subset of MEC neurons showed strikingly periodic activation as a function of distance run (blue), whereas others did not (black). **C)** Autocorrelation of the entire firing rate trace for the 6 neurons shown in B. To identify distance cells, we compared autocorrelation peaks to within-cell shuffled distributions. **D)** Peak prominence versus peak z-score (normalized to shuffled distribution) for all 14,172 neurons recorded in dark sessions. Distance cells (blue) were defined as those with peak prominence > 0.1 and shuffle p-value < 0.01. **E)** Heatmaps of the firing rate autocorrelation for all distance cells in four example sessions, with cells sorted from dorsal (top) to ventral (bottom). The modularity and depth-dependence of distance tuning, reminiscent of grid cells, is evident. **F)** Number and percentage of neurons identified as distance-tuned in all 57 sessions, colored by mouse (number distance-tuned cells, mean ± SEM = 28 ± 3, range = 0 – 111; percentage, mean ± SEM over sessions = 13 ± 2%, range = 0 – 52%). **G)** 2D histogram of number of distance cells versus preferred distance (period) and distance from MEC entry. Most distance cells were found in the most dorsal 1000 μm of MEC. A substantial fraction of distance cells were found up to 400 μm above the border of MEC, but the fraction dropped sharply after that, indicating a possible ventral bias in the precise identification of the MEC boundary. **H)** Histogram of distance cell periods (marginal histogram of G; 2 cm bins, smoothed with a Gaussian kernel with standard deviation 1 bin). Dashed lines indicate 75cm times powers of 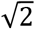. **I)** Same as H, but with individual mice indicated by different colors. **J)** Average shift (computed as location of the peak cross correlation) for distance cells versus non-distance cells during gain = 0.8 Cluster 1 (stable) blocks. To be included in this analysis, cells had to pass the same spatial stability criteria used in the rest of the manuscript (stability score ≥ 0.5 in 6 baseline trials). Map shifts were significantly larger (more negative) for distance cells compared to non-distance cells (mean map shift ± SEM, distance cells = −9.2 ± 0.5 cm, N = 163 cells from 8 sessions in 5 mice; non-distance cells = −5.2 ± 0.2 cm, N = 599 cells from 12 sessions in 8 mice; p = 3.8e-16, Mann-Whitney U test). **K)** Difference in average shift between distance and non-distance cells for each gain change block that had at least 5 of each (difference, distance cells – non-distance cells = −2.7 ± 0.6 cm, N = 8 gain change blocks from 6 sessions in 4 mice, p = 0.016, Wilcoxon signed rank test). Different colors correspond to different mice. **L)** Trial-to-trial correlation matrices for distance cells and non-distance cells in remapping blocks (clusters 2 and 3). **M)** Single trial peak correlation to the average baseline firing rate map across the gain change block, separated by distance and non-distance cells. Remapping was more pronounced in distance cells compared to non-distance cells (dots indicates p < 0.001, no dots indicates not significant, Mann-Whitney U test on distance cells vs. non-distance cells for each trial). **N)** Optimal delay factors, computed as in Fig. 4D, for distance and non-distance cells separately. Distance cells’ delay factors were significantly more negative (prospective) than for non-distance cells (delay factor, mean ± SEM, distance cells = −49.3 ± 3.8 ms, N = 549; non-distance cells = −13.2 ± 2.0 ms, N = 2740; p = 6.2e-14, Mann-Whitney U test). **O)** Same as N, averaged by session (difference in temporal delay, distance cells – non-distance cells, mean ± SEM over sessions = −36.5 ± 6.2 ms, N = 23 sessions from 7 mice, p = 3.5e-5, Wilcoxon signed rank test). Different colors correspond to different mice. **P)** Change in map shift in low contrast (10% and 20%) compared to high contrast (100%), a signature of Kalman filtering. Changes in map shift were larger (more negative) for distance cells compared to non-distance cells (mean ± SEM, distance cells, c=10: −1.8 ± 1.8 cm, N = 52; c=20: −2.3 ± 0.9 cm, N = 97; non-distance cells, c=10: −0.5 ± 0.8, N = 138; c=20: −0.2 ± 0.5 cm, N = 317; p = 0.034 for difference between distance and non-distance cells, p = 0.90 for difference between c=10 and c=20, n-way ANOVA). Boxes indicate the 25^th^, 50^th^, and 75^th^ percentiles, whiskers indicate the range of data points not determined to be outliers, and circles indicate outlier data points. **Q)** Cumulative percent variance explained by principal components of activity of distance and non-distance cells during running in the dark (Methods). For this analysis, we chose sessions with at least 30 distance and non-distance neurons (N = 15 sessions from 7 mice). For each such session, we randomly selected 30 distance and non-distance neurons and performed PCA on each sub-sampled population. Small dots above the plot indicate significance of Wilcoxon signed rank tests between distance cells and non-distance cells (three dots, p < 0.001; two, p < 0.01; one, p < 0.05; no correction for multiple comparisons).

As previous work demonstrated that MEC grid cells respond more to VR gain changes than border cells (Campbell et al., 2018), we asked whether VR gain changes impacted distance cells (putative grid cells) differently than other MEC neurons. In Cluster 1 blocks (no remapping, Figure 2), distance cells’ spatial maps shifted by more than non-distance cells (Figure 6J, K), consistent with a larger influence of path integration. These results demonstrate that a distribution of path integration responsiveness exists in MEC, with putative grid cells occupying the tail of this distribution. The presence of a distribution of path integration responsiveness is consistent with the observation that MEC map shifts were correlated across multiple gain change repetitions (Figure S3C,D).

We next asked whether distance cells responded differently than other MEC neurons during gain change-induced remapping (Clusters 2 and 3, Figure 2). Indeed, although distance cells had higher baseline stability than non-distance cells in the 6 trials preceding the gain change, the peak correlation to baseline trials following gain change onset dropped by more in distance cells than non-distance cells (Figure 6L, M). This indicates that distance cells (putative grid cells) remapped more in response to gain changes than other MEC neurons, consistent with previous work in which grid cells were identified using electrophysiological recordings in freely moving mice foraging in an open field (Campbell et al., 2018).

We hypothesized that MEC uses path integration calculations to predict position tens of milliseconds into the future, prior to the arrival of delayed sensory input (Figure 4E). If dark distance cells drive path integration calculations in MEC, then dark distance cells should show an even more pronounced prospective bias relative to other MEC cells. Indeed, the optimal temporal delay factor was −49.3 ± 3.8 ms for distance cells versus −13.2 ± 2.0 ms for non-distance cells (p = 6.2e-14, Mann-Whitney U test; Figure 6N, O). Moreover, consistent with the idea that Kalman filtering could be implemented by grid cell attractor dynamics, signatures of Kalman filtering were more pronounced in distance cells than non-distance cells. Specifically, changes in map shift in low contrast were larger for distance cells than non-distance cells (Figure 6P).

Finally, attractor dynamics in MEC have been suggested to force grid cell activity to occupy a low-dimensional manifold (Yoon et al., 2013). If this is true, then we would expect the dimensionality of distance cell (putative grid cell) activity to be lower than the dimensionality of non-distance cell activity. To check this, we performed PCA on the activity traces of matched numbers of distance and non-distance cells in each dark session that had at least 30 of each. A greater percentage of variance was explained by low dimensional PC spaces for distance cells compared to non-distance cells (Figure 6Q), even when cells were matched for mean firing rate or depth within the MEC (Figure S6). These results support the hypothesis that the activity of putative MEC grid cells occupies a low dimensional manifold. The ability to record many such cells simultaneously (max 111 here) will open avenues for investigation into the geometry of this low dimensional activity space.

### Large gain changes reveal stronger influence of non-visual signals in RSC compared to V1

As previous work indicated that large visuomotor cue conflicts reliably trigger remapping in MEC (Campbell et al., 2018), we next considered how large conflicts between landmarks and path integration cues in the current VR track influence MEC, V1 and RSC. For these experiments, we reduced the gain between the rotation of the running wheel and the mouse’s progression along the virtual hallway by a factor of 0.5 for blocks of 4 trials (number of blocks per session = 1-4, median 2) (Figure 7A, B). As expected, the gain = 0.5 condition reliably caused remapping in MEC (Figure 7C, left), whereas V1 neurons remained locked to visual cues (Figure 7C, middle). However, unlike the gain = 0.8 condition, in the gain = condition the spatial firing fields of RSC neurons also showed signatures of remapping and often shifted, appeared, disappeared or changed in firing rate (Figure 7C, right).

**Figure 7:**
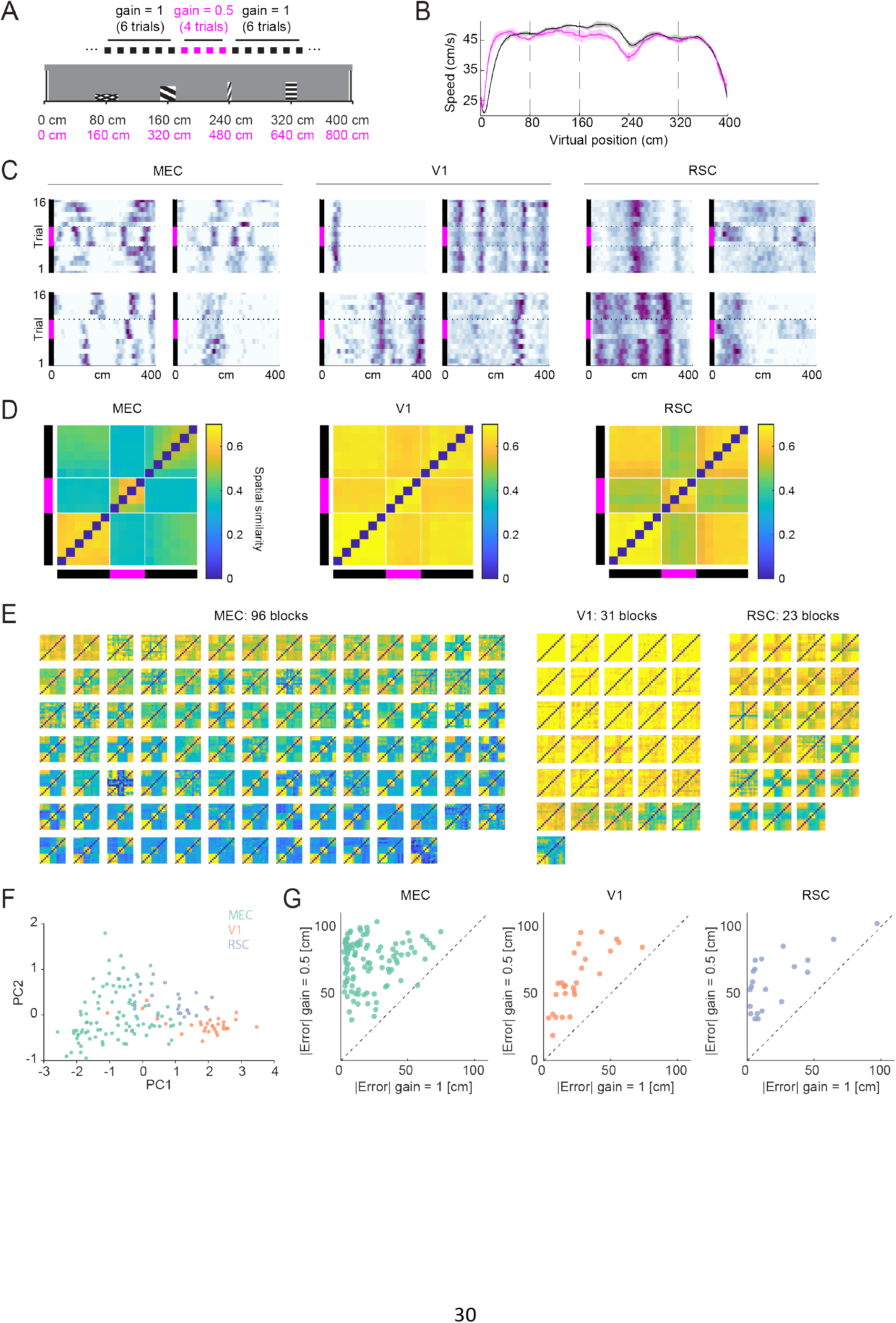
Large gain changes reveal an influence of non-visual cues in RSC. **A)** Illustration of the large gain change experiment design. Blocks of four trials in which the VR gain was set to 0.5 were surrounded by blocks of six trials in which gain was 1 (same design as g=0.8 experiment, Figure 2A). **B)** Average running speed versus track position (solid lines) ± SEM (shaded area) over blocks (N = 208 blocks from 30 mice) during baseline trials (black) and g=0.5 trials (magenta). Average running speed did not differ between the conditions (p=0.33, Wilcoxon signed rank test). Dashed lines indicate tower locations. **C)** Spatial firing rate maps of example neurons in MEC (left), V1 (middle), and RSC (right) during large gain change blocks. All twelve neurons are from different mice. Black bars indicate baseline trials, magenta bar indicates trials with g=0.5. Most MEC neurons remapped, although there were some exceptions (e.g. example MEC cell on the bottom right). In contrast, almost no V1 neurons remapped. Unlike for the g=0.8 condition (Figure 2C), RSC neurons often remapped in response to g=0.5. **D)** Average similarity matrix by brain region for all large gain change blocks (MEC: 96 gain change blocks, 56 sessions, 18 mice, median 2 [max 4] gain change blocks per session, 41 ± 4 [SEM] spatial cells per block; V1: 31 blocks, 18 sessions, 6 mice, median 2 [max 2] gain change blocks per session, 22 ± 2 [SEM] spatial cells per block; RSC: 23 blocks, 12 sessions, 7 mice, median 2 [max 2] gain change blocks per session, 50 ± 7 [SEM] spatial cells per block). Similarity decreased sharply at the onset of gain change for MEC, and to a lesser extent, RSC. Similarity remained high during g=0.5 in V1. **E)** Average similarity matrices as in (D), but for individual gain change blocks. **F)** The first two principal components of the trial-trial similarity matrices shown in (E). Each point represents an individual gain change block. V1 blocks were highly separable from MEC blocks, and RSC blocks were intermediate between the two. **G)** Average absolute decoder error in baseline trials versus gain change trials in MEC, V1, and RSC. The decoder was trained on baseline trials, which resulted in consistently higher errors in gain change trials. Points near the y-axis indicate remapping (low decoder error in baseline, high decoder error during gain change) and can be seen most prominently in MEC but also in RSC. Baseline decoder errors were larger in V1 due to tower misclassifications (Figure 1K, Figure S1C).

These neuron-level observations were confirmed by population-level analyses (Figure 7D, E, F). We selected spatial cells in the same manner as for g=0.8 trial blocks and computed average trial-trial peak cross-correlation matrices over trial blocks with at least 5 spatial cells. These matrices revealed large changes in spatial maps in MEC, intermediate changes in RSC, and almost no change in V1 (Figure 7D). Examining average trial-trial correlation maps for individual trial blocks revealed that MEC populations almost exclusively remapped in response to g=0.5, whereas V1 populations remained locked to landmarks, and RSC showed varying degrees of remapping (Figure 7E). PCA analysis demonstrated that RSC trial blocks were intermediate between V1 and MEC (Figure 7F). A complementary decoding analysis gave similar results (Figure 7G). Thus, creating larger conflicts between path integration and landmark cues revealed an influence of non-visual inputs on RSC spatial maps that was not apparent during smaller cue conflicts (Figure 2). However, remapping in RSC was qualitatively different than remapping in MEC. Specifically, RSC maps always returned to their pre-gain change state in the baseline trials following the gain change, whereas this was not necessarily the case in MEC (Figure 7D, E, Figure S7B).

Finally, we sought to determine the temporal dynamics of remapping in MEC. Remapping occurred shortly after gain change onset (Figure S7D, median 3.9s, 2.5s 6.0s 25^th^ and 75^th^ percentile). Prior to remapping, the decoding error tended to steadily increase (Figure S7E). In a previously described coupled oscillator attractor model of gain change-induced remapping (Campbell et al., 2018; Ocko et al., 2018), the phase shift between the landmark input and path integration attractor steadily increases after the onset of gain change until a decoherence threshold is reached (Figure S7F). Thus, the temporal dynamics of MEC remapping were qualitatively consistent with this coupled oscillator model.

## Discussion

Here, we examined the degree to which neurons in three cortical regions involved in navigation followed unique versus shared coding principles. While recent work has revealed distributed and heterogeneous coding across cortical regions involved in navigation (Allen et al., 2019; Clancy et al., 2019; Hardcastle et al., 2017; Minderer et al., 2019; Musall et al., 2019; Pinto et al., 2019; Stringer et al., 2019), we report that MEC, V1 and RSC implement distinct cue integration algorithms during navigation. Path integration influenced position estimates in MEC more than in V1 and RSC. Path integration calculations could allow MEC to predict an animal’s current or future position before the arrival of delayed visual input. Indeed, MEC neurons coded position with a small prospective bias (tens of milliseconds). In MEC, but not V1 or RSC, path integration position estimates combined with landmarks following the principles of a Kalman filter, which can be approximately implemented via attractor dynamics. On the other hand, V1 and RSC were strongly influenced by visual landmark features, with delays in sensory processing modulating spike positions in both areas (retrospective coding). However, large conflicts between path integration and landmarks revealed an influence of path integration in RSC (gain change-induced remapping) that was not present in V1. Together, this work suggests that during navigation, cortical regions implement distinct algorithms for combining path integration with landmarks, which could allow specialization in how cortical brain regions optimally support different navigational strategies.

Other works have emphasized the influence of non-visual signals, including path integration and visuomotor predictions, on activity in mouse V1 (Fiser et al., 2016; Fournier et al., 2020; Guitchounts et al., 2020; Keller et al., 2012; Saleem et al., 2018). Consistent with this work, our data reveal an influence of path integration in V1, in that maps shifted during gain changes by an amount that could not fully explained by delays in sensory processing. However, the influence of path integration was much stronger in MEC than V1, and we could not induce remapping in V1 even for large gain changes that induced remapping in RSC. Thus, although path integration can exert a small influence on V1 spatial representations, V1 maps remain locked to visual cues in our task even when RSC and MEC remap.

Our findings also have implications for how neural representations in RSC uniquely support navigation. We found visual landmarks strongly influenced RSC neural firing patterns, consistent with work demonstrating that many RSC neurons encode visual landmarks (Fischer et al., 2020). On the other hand, path integration had less influence on RSC compared to MEC neural representations in the current VR task. This contrasts with previous work demonstrating that lesioning or inactivating RSC impairs path integration in darkness (Cooper and Mizumori, 1999; Elduayen and Save, 2014) and that RSC shows degraded neural responses when animals are passively shown the visual stimuli associated with a VR environment (Fischer et al., 2020). However, task engagement and locomotion-driven changes in brain state can alter RSC neural representations (Fischer et al., 2020) and correlations with other brain regions (Clancy et al., 2019). In the current work, conditions where path integration and locomotor conflicts were large revealed an influence of non-visual inputs on RSC spatial maps. One possibility is that contextual changes such as our large gain change could alter RSC’s functional connectivity, upweighting inputs from the hippocampal formation and triggering remapping. In contrast, during small gain changes, when path integration and landmarks do not disagree by much, input from visual cortex could dominate over feedback from the hippocampal formation. This flexibility could allow the mouse to make use of different types of spatial representations in different contexts. Thus, while the current VR task offers a window into comparing the default cue integration algorithms across MEC, RSC and V1, future work should examine the rigidity versus flexibility of these cue integration algorithms under changing task demands or behavioral states.

Despite the advantages afforded by head-fixation and virtual reality, it remains challenging to study MEC coding properties in head-fixed mice as 1D grid cell firing patterns are difficult to distinguish from those of non-grid MEC cells including border cells, cue cells, and irregular spatial cells (Campbell et al., 2018; Domnisoru et al., 2013; Kinkhabwala et al., 2020). However, we observed a population of MEC neurons tuned to distance run in the dark (~11% of MEC cells, range 0-52%) in which MEC’s unique cue integration properties were most pronounced. The striking modularity of distance tuning in this cell population supports the idea that these neurons correspond to MEC grid cells. Other properties of distance tuned cells were also consistent with grid cells: compared to other MEC neurons they were more influenced by path integration (Campbell et al., 2018), showed increased prospective temporal delay (Kropff et al., 2015), and their population activity showed a lower-dimensional structure (Yoon et al., 2013). The possibility that grid cells can be identified in 1D head-fixed VR environments simply by allowing mice to freely run in the dark would improve the field’s ability to bring head-fixed techniques to bear to dissect MEC neural circuits.

The majority of electrophysiological studies on MEC to date have considered, at most, the simultaneous activity of tens of cells and thus, characterizing variability in MEC neural representations at the level of single sessions and animals has remained challenging. By simultaneously recording from an average of 166 MEC cells per recording session, we shed light on the consistency of MEC neural responses at the single trial, session and animal level. We find that, at the population level, variability in MEC neural responses can clearly be observed across animals, sessions and trials, as well as at the single cell level within a given session. In particular, we observed pronounced variability in how MEC neurons weighed visual landmark versus locomotor cues. These results suggest a distribution of path integration responsiveness exists in MEC, with putative grid cells occupying the tail of this distribution. This differs from V1, in which visual landmarks dominated position coding in nearly every recording session, and from RSC in which the influence of path integration was only revealed under conditions in which cue conflicts were very large. Understanding how cue integration and representations are transformed across these regions is of interest for future work.

Our previous work proposed a coupled oscillator attractor model to explain MEC grid cell responses to gain manipulations, specifically the sharp transition between coherent shifts and remapping (Campbell et al., 2018; Ocko et al., 2018). We show that for small cue conflicts, the attractor model’s position estimates closely resemble the output of a Kalman filter (Deneve et al., 2007; Wilson and Finkel, 2009). Gain change experiments in low contrast conditions confirmed the dependence of map shifts on the uncertainty of landmark input, consistent with both models. Thus, we hypothesize that attractor dynamics form the mechanistic basis of Kalman filter-like computations in MEC. However, the attractor model as it stands has no representation of uncertainty, which is one of the hallmarks of a Kalman filter. One possibility is that uncertainty is represented in amplitude of the attractor bump or the variability of the bump location. Moreover, whereas the Kalman model can be modified to represent predicted future position by adding an additional path integration-based calculation, the attractor model represents exact current position, which is inconsistent with the prospective bias we observed in putative MEC grid cells. On the other hand, a Kalman filter fails to capture the transition between coherent shifts and remapping. Future work should explore how attractor models can be altered to incorporate representations of uncertainty and make short time scale predictions of future position.

Further work is also needed to understand the relationship (if any) between the short time scale prospective coding observed here (tens of milliseconds into the future), which was most pronounced in putative MEC grid cells, and the longer time scale predictions that could create successor-like representations in the hippocampus and MEC to support reinforcement learning (Stachenfeld et al., 2017). While these two types of predictions operate on very different time scales, it is possible that mechanisms for making short time scale predictions based on path integration could be co-opted to make longer time scale predictions about the occupancy of non-local states.

## Acknowledgments

We would like to thank Bob Schneeveis for engineering assistance, Adriana Diaz for histology assistance, and members of the Giocomo laboratory for feedback and discussions.

## Author Contributions

MGC, AA, and LMG conceived the project and designed the experiments. MGC and AA collected and analyzed data. SAO performed modeling and theoretical analysis under supervision of SG. MGC, AA, and LMG wrote the paper with feedback from SAO and SG.

---

**Figure S1:**
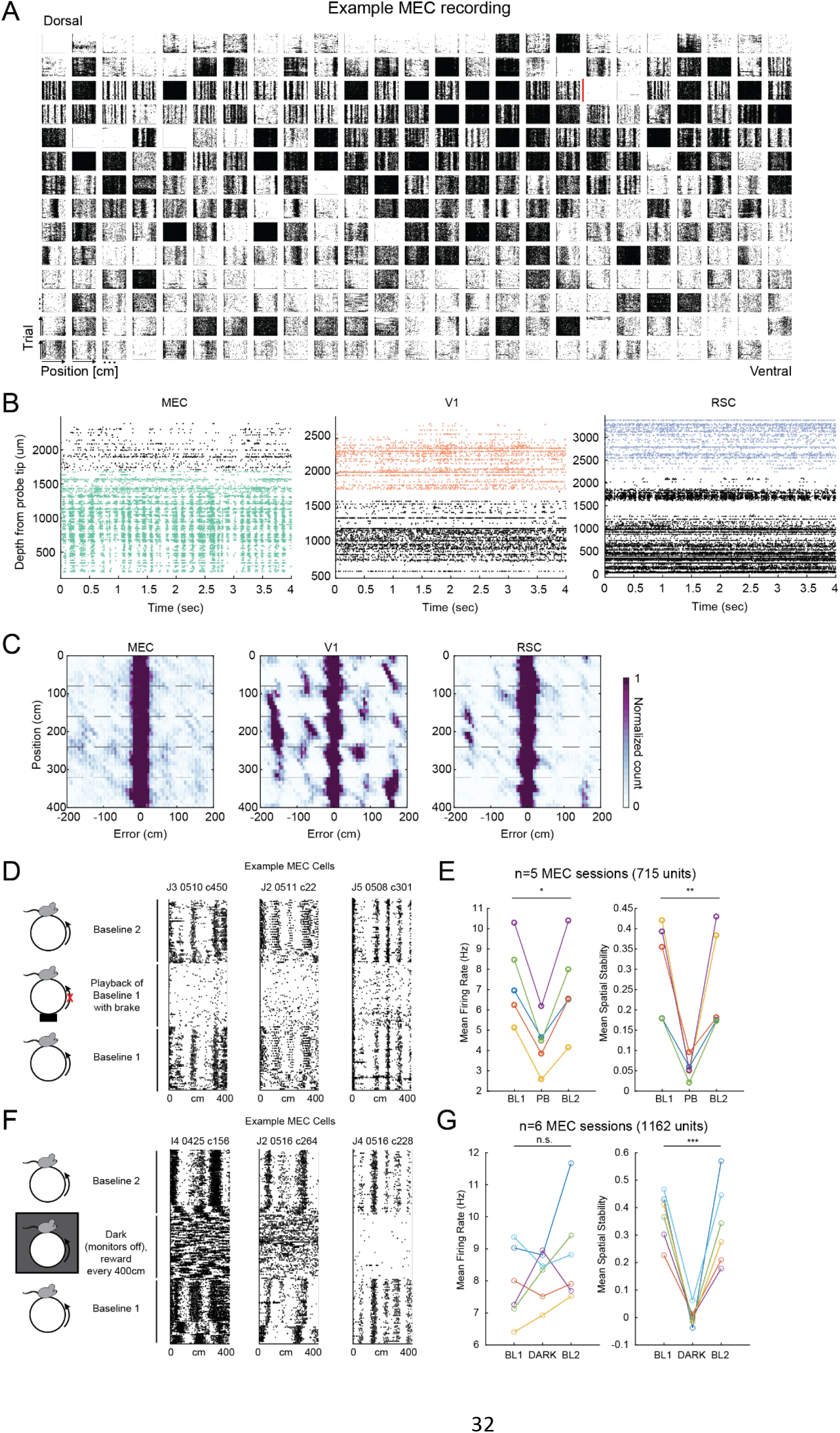
Raster plots of MEC neurons, theta-paced traveling waves in MEC, and the influence of active behavior and darkness on MEC firing patterns. **A)** Raster plots of 350 simultaneously recorded neurons from a session targeting MEC. Each subpanel shows the raster plot of a single neuron along the virtual reality track (x-axis) for 50 baseline trials (y axis). Red line shows the boundary of MEC estimated from histology. Plots ordered according to the dorsal-ventral position of the neuron, from left to right and then top to bottom. **B)** Raster plots of all recorded spikes over example 4 second snippets ordered by distance from probe tip (y-axis). Spikes are colored if they were identified as within the target brain region (green: MEC, red: V1, blue: RSC), and black otherwise. Consistent with previous work (Hernández-Pérez et al., 2020), MEC neurons activated in a theta-paced wave that traveled from dorsal to ventral MEC (left). This activity pattern was not observed in V1 or RSC (center, right). **C)** Decoding error map shows how the distribution of decoding error varies along the track for the decoder used in Figure 1K. For a decoder trained on MEC neurons (left), the error is distributed evenly around 0. In contrast, for a decoder trained on V1 neurons (middle), and to a lesser extent for a decoder trained on RSC neurons (right), a higher frequency of errors of around 80cm or 160cm indicates misclassification of towers. **D) Left**: Experiment structure for passive replay (playback) experiment. In a subset of mice (n=4), we investigated the response properties of MEC neurons as mice passively viewed traversals of the virtual corridor. In these experiments, mice first actively traversed the hallway for 50 trials (baseline 1). We then prevented mice from running by blocking the motion of the running wheel and replayed the self-generated traversals from baseline 1. Mice therefore passively viewed the identical visual stimulus (playback). Note that these mice were habituated to the blocking of the running wheel already during training (Methods). Playback sessions were followed by 50 trials of a second baseline session (baseline 2). **Right**: Raster plots of three example MEC units for baseline 1, playback and baseline 2 sessions. **E)** Average firing rates (left) and spatial stability (right, calculated as the correlation between the firing rate map computed in the first half vs second half of trials, within 10 trial blocks, averaged over trial blocks) were both reduced during the playback session, highlighting the importance of active behavior for spatial firing in MEC (mean firing rate ± SEM, BL1 = 7.4 ± 0.9 Hz, PB = 4.3 ± 0.6 Hz, BL2 = 7.1 ± 1.0 Hz, ANOVA F(2,12) = 3.9, p = 0.0496; spatial stability ± SEM, BL1 = 0.30 ± 0.05, PB = 0.06 ± 0.01, BL2 = 0.27 ± 0.06, ANOVA F(2,12) = 8.9, p = 0.0043; n = 5 sessions from 4 mice). *** p < 0.001, ** p < 0.01, * p < 0.05, n.s. not significant BL1: Baseline 1, PB: Playback, BL2: Baseline 2. **F) Left:**Experiment structure for dark sessions with reward. In a subset of mice (n=5), we investigated the response properties of MEC neurons when mice were running in darkness, but still receiving reward every 400 cm. **Right:** Raster plots of three example MEC units during baseline 1 (BL1), dark sessions (DARK), and baseline 2 (BL2) (trials per condition, left: 70, middle: 50, right: 50). The total number of trials in each condition was 50-220 (median 75) for BL1, 50-130 (median 50) for DARK, and 50-70 (median 50) for BL2 (n=6 sessions from 5 mice). **G) Left:** Average firing rate for baseline and dark blocks. Although we observed a variety of effects across cells (see examples), firing rates did not change on average (mean firing rate ± SEM, BL1 = 7.9 ± 0.5 Hz, DARK = 8.2 ± 0.3 Hz, BL2 = 8.8 ± 0.6 Hz, n = 6 sessions from 5 mice, ANOVA F(2,15) = 1.0, p = 0.39). **Right:** Low average spatial stability during darkness emphasizes the importance of visual cues for spatial firing in MEC (spatial stability ± SEM, BL1 = 0.37 ± 0.04, DARK = 0.00 ± 0.01, BL2 = 0.34 ± 0.06, n = 6 sessions from 5 mice, ANOVA F(2,15) = 23.4, p = 2.4e-5). *** p < 0.001, ** p < 0.01, * p < 0.05, n.s. not significant.

**Figure S2:**
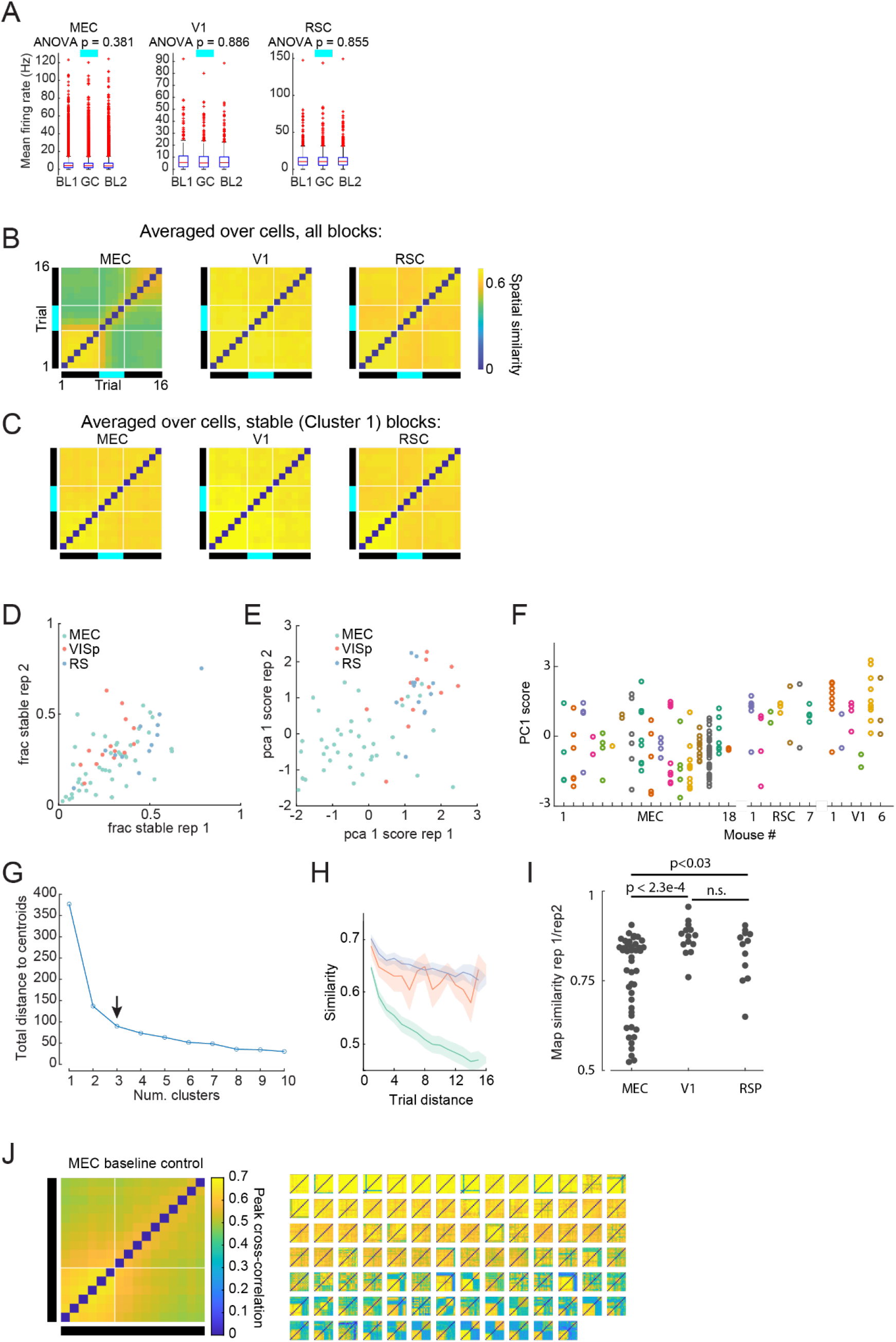
Additional characterization of responses to the gain=0.8 condition and baseline controls. **A)** Firing rate of neurons was similar between baseline and gain change trials in all brain regions. Only spatial cells (stability score ≥ 0.5 in baseline trials) were included in this analysis, but results were similar for all neurons (data not shown). Boxes indicate 25^th^ and 75^th^ percentiles, red line indicates median, whiskers indicate range of data points not determined to be outliers, red dots indicate outliers. **B)** Spatial similarity maps averaged across all spatial neurons in MEC (n=6008 of 23868 cells, left), V1 (n=571 of 1881 cells) and RSC (n=1373 of 3436 cells) for gain=0.8 blocks. Note the increased similarity between pre baseline, gain trials and post baseline in V1 and RSC neurons compared to MEC neurons, consistent with the observation of increased remapping during gain trials in MEC, but not V1 and RSC. **C)** Same as (B), but for spatial neurons from stable blocks (cluster 1) only (MEC: 1709 of 5412 units, V1: 531 of 1662 units, RSC: 1315 of 3218). **D)** Comparison of the fraction of stable neurons across pre-baseline in sessions with two blocks of gain=0.8. There was a slight decrease in the fraction of stable units in MEC and RSC, and no change in V1 (one-way ANOVA F(2,67)=4.51, p<0.015, decrease in MEC p=0.01, 44 pairs, decrease in RSC p=0.007, 12 pairs, no change in V1, p=0.152, 14 pairs, Wilcoxon signed rank test). **E)** Same as in D), but for the score of the first principal component (PC1). **F)** Distribution of PC1 score for each brain region, separated by mouse. Each dot represents a gain change block and colors correspond to different mice. In MEC, most mice showed a wide variety of PC1 scores, indicating different types of responses during gain = 0.8 trials. G) Total distance to centroid in PCA space for different numbers of clusters in clustering gain change responses using k-means (as in Figure 2F). Arrow indicates the chosen number of clusters (k=3), which lay at the elbow of this curve. **H)** Pairwise spatial similarity as a function of inter trial distance (ITD) for blocks of 16 baseline trials. While pairwise spatial similarity was similar for V1 and RSC (Stability ITD 1, no difference between V1 and RSC, p>0.85; stability ITD 15 no difference between V1 and RSC, p>0.55), pairwise similarity was lower and decreased more in MEC compared to V1 and RSC (decrease between stability IDT 1 and IDT 15, one-way ANOVA f(2,115)=10.63, p<5.8e-5, decrease for MEC larger than for V1, p<1.5e-4, decrease for MEC larger than RSC, p<0.016, no difference in decrease for V1 and RSC, p>0.38). This reflects the fact that MEC maps often spontaneously remapped. All tests Mann-Whitney U tests. Shading indicates s.e.m. **I)** Spatial similarity between the average spatial firing rate for the 6 trials before the first gain change onset and second gain change onset in sessions with two or more gain=0.8 blocks (MEC: 44 pairs, V1: 14 pairs, RSC: 12 pairs). Stability in MEC is lower than V1 and RSC (one-way ANOVA, f(2,67)=7.68, p<0.001, SimlarityMEC<SimilarityV1: p<2.3e-4, SimilarityMEC<SimilarityRSC: p<0.033, Mann-Whitney U test). **J)** Left: Average spatial similarity map for blocks of 16 baseline trials for MEC recordings (96 blocks from 53 sessions in 17 mice). Right: Average similarity maps for individual blocks of baseline trials, sorted by their projection onto principal component 1 from gain change blocks (see Methods), showing that spontaneous remapping occurs in baseline trials.

**Figure S3:**
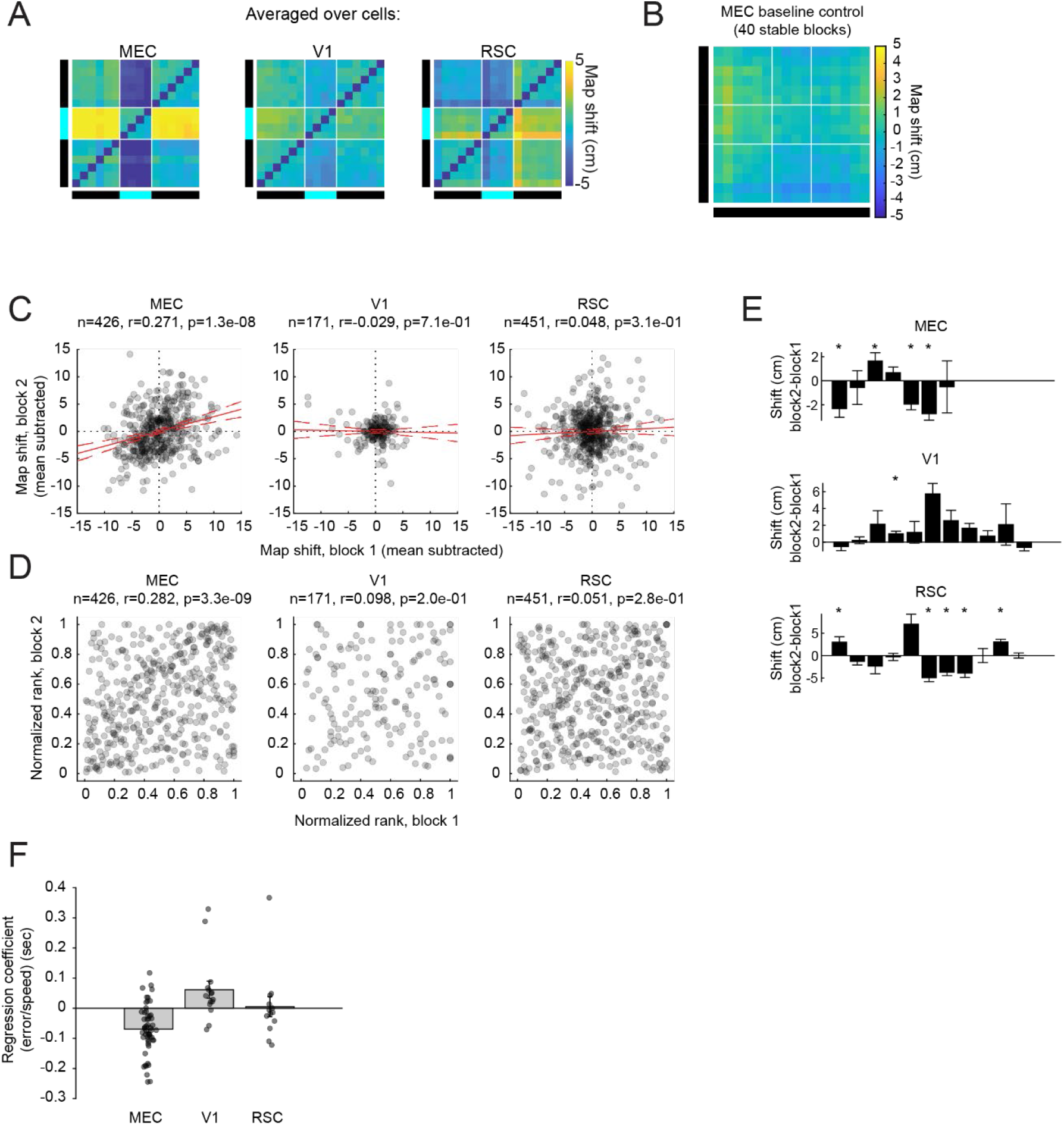
Additional characterization of map shifts during the gain=0.8 condition and decoding-based analysis of prospective versus retrospective coding. **A)** Trial by trial shift for gain=0.8 blocks, averaged across spatial neurons from stable blocks only (MEC: 1709 of 5412 units, V1: 531 of 1662 units, RSC: 1315 of 3218 units). Black bars indicate baseline trials and cyan bar indicates gain trials. **B)** Same as Figure 3C but for blocks of 16 baseline trials in MEC recordings. There was a small non-zero map shift for MEC baseline recordings (−0.9 ± 0.3 cm, N = 40 blocks from 27 sessions in 14 mice; p = 0.012 versus zero shift, Wilcoxon signed rank test; shift computed for each block as in Figure 3D), possibly due slow drifts or asymmetric expansions of firing fields over time (Mehta et al., 2000). However, these shifts were significantly smaller than map shifts during the gain = 0.8 condition (−4.1 ± 0.5 cm, Figure 3D, p = 1.5e-5, Mann-Whitney U test). **C)** Map shift of individual neurons in block 1 on x axis (subtracted by mean across all simultaneously recorded spatial cells from block 1), and map shift of individual neurons in block 2 on y axis, for sessions with two or more gain = 0.8 repetitions, where both blocks were classified as ‘stable’ (Cluster 1). In MEC, but not V1 and RSC, map shifts are significantly correlated across blocks. **D)** Relative order of the shift size for gain change block one and two for same neurons as in (C). We determined the ranking based on the map shift of all simultaneously recorded spatial cells within each gain change block and compared ranks across blocks. This controls for the possibility that correlations in (C) were driven by session-level differences in the mean map shift. In MEC, but not V1 and RSC, neurons that shifted most in block 1, also tended to shift more in block 2. **E)** Same data as in (D), but split by into individual pairs of reps (MEC: 7 pairs of Cluster 1 reps from 5 sessions in 5 mice; V1: 11 pairs of reps from 11 sessions in 4 mice; RSC: 11 pairs of reps from 11 sessions in 7 mice). Bars represent average difference in shift between rep 1 and rep 2, and error bars indicate standard error of the mean over cells. Stars indicate sessions with a significant difference between the mean map shift in rep 1 versus rep 2 (p < 0.05, Wilcoxon signed rank test). **F)** Results of decoding approach to estimating optimal delay (related to Figure 4). Position was decoded from population activity in each brain region, and decoding errors were regressed on running speed and position using multiple linear regression (Methods). The slope of the regression coefficient on running speed, shown here, is an estimate of the degree of retrospective (positive coefficients) versus prospective (negative coefficients) coding. In qualitative agreement with Figure 4E, V1 coded position retrospectively, whereas MEC coded position prospectively, and RSC fell between the two extremes (mean coefficient ± SEM [sec], MEC = −0.069 ± 0.011, N = 54 sessions; V1 = 0.062 ± 0.028, N = 15 sessions; RSC = 0.005 ± 0.033, N = 13 sessions; ANOVA F(2,79) = 12.96, p = 1.4e-5).

**Figure S4:**
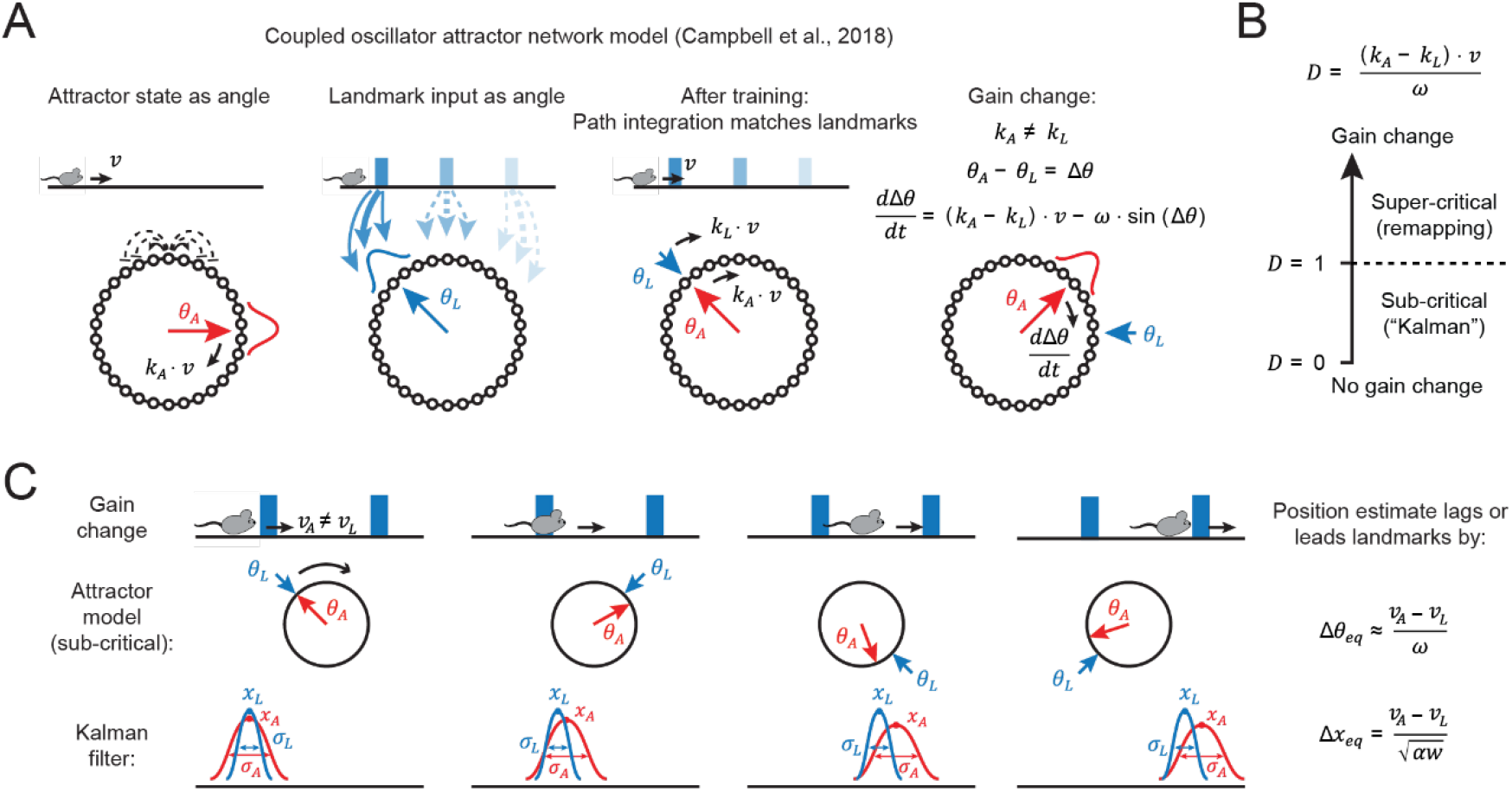
An attractor network model recapitulates the output of a Kalman filter in response to small gain changes. **A)** Cartoon illustrating the grid cell attractor network model from Campbell et al., 2018. Left: The ring attractor architecture in which neurons excite their nearest neighbors and inhibit their next-nearest neighbors. This leads to a set of steady state activity patterns corresponding to unimodal “bumps” of activity on the ring. The location of the bump on the ring can be parameterized by an angle, *θ_A_*. Adding conjunctive position-velocity inputs into the network allows it to perform path integration, in that the activity bump moves around the ring at a speed equal to *k_A_* ⋅ *v*, where *v* is the running speed of the mouse and *k_A_* is a constant describing the grid cell spatial frequency. Middle: Landmark inputs reinforce the correct phase relative to the external landmarks. The location of the input relative to the attractor ring can be parameterized as angle, *θ_L_*. Right: After training, path integration matches landmarks, such that that *θ_A_* and *θ_L_* move around the ring at the same rate. During a gain change, the difference between the attractor phase, *θ_A_*, and the landmark phase, *θ_L_*, evolves according to the differential equation shown at top. **B)** Illustration of the boundary between the sub-critical (coherent map shifts) and super-critical (remapping) regimes of the attractor model. The transition occurs when the decoherence number, *D*, equals 1. **C)** Cartoon illustrating the correspondence between the sub-critical regime of the attractor model and the output of a Kalman filter. During a gain change, the animal’s estimated running speed, *v_A_*, does not equal the speed of the landmarks, *v_L_*. The cartoon illustrates a situation where *v_A_* > *v_L_* (gain decrease, as in the experiments in this paper). In this situation, both models arrive at an equilibrium map shift in which the estimated position is ahead of the true position. In both models, the map shift is directly proportional to the difference between *v_A_* and *v_L_* and inversely proportional to a term reflecting landmark strength. In the Kalman filter model, this term is 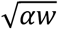, where *α* is a diffusivity constant that describes how position uncertainty grows over time during path integration, and *α* is the “Kalman landmark strength” in the continuous limit, reflecting the strength, or certainty, of landmark input (Methods). The analogous quantity to 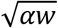 in the attractor model is *w* (the strength of the landmark input).

**Figure S5:**
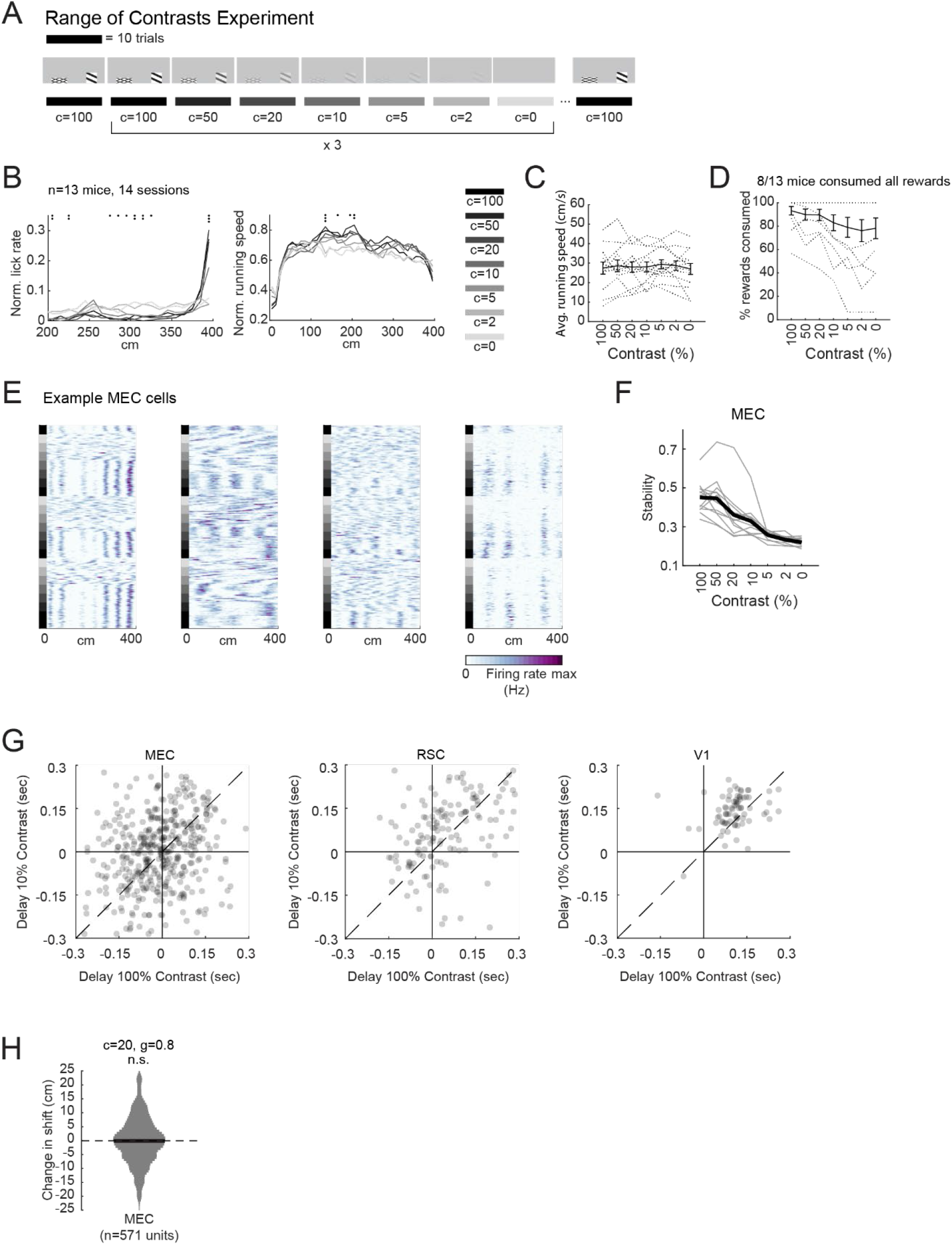
Lowering visual contrast reduces the precision of anticipatory licking and reduces the stability of MEC spatial maps. **A)** Experimental design to test the influence of visual contrast on spatial firing rate maps. Contrast (c) was reduced from 100 down to 0 in 10 trial blocks. The entire sequence was repeated 3 times. **B)** Lick rate (left) and running speed (right) as a function of VR position for different contrast levels. For high contrast levels, mice tended to lick only close to the reward tower, for lower contrast levels, licking became more distributed along the track, indicating that the mice were less certain about their location within the VR environment (three dots, p < 0.001, two dots, p < 0.01, one dot, p < 0.05, one-way ANOVA). **C)** Average running speed did not depend on contrast (one-way ANOVA, F(6,84) = 0.09, p = 0.997). Black line shows means, error bars SEM, dotted lines individual mice. **D)** Percent rewards consumed as a function of contrast. Eight out of 13 mice consumed all rewards at all contrast levels. The other 5 mice consumed fewer rewards as contrast was reduced. Black line shows means, error bars SEM, dotted lines individual mice. **E)** Example MEC neurons’ firing rate maps during the range of contrast experiment. Squares on the left of the plot indicate contrast level, as in panel (A). **F)** Spatial stability decreases with decreasing levels of contrast. In a subset of mice (13 sessions, 12 mice), we recorded activity in MEC while successively decreasing the contrast level of the visual landmarks. We found that spatial stability decreased below baseline level for c=20 and lower (c_50_: p>0.89, c20: p<0.002, c10-c0, all p<2.4e-4, Wilcoxon signed rank test). **G)** Estimated temporal delay during baseline (c=100) and low contrast (c=10). Lowering the contrast did not affect the estimated temporal delay in MEC (417 units from 17 sessions, p>0.07), but increased temporal delay in RSC (136 units from 8 sessions, p<0.008) and in V1 (78 units from 7 sessions, p<4.1e-5). Each dot represents a neuron. All tests Wilcoxon signed rank test. **H)** In a subset of mice in which we targeted MEC for recording, we performed gain=0.8 experiments in contrast = 100 (c=100) and contrast = 20 (c=20). The analysis is identical to Figure 5F. There was no significant difference in map shift between c=100 and c=20 (change in map shift, c20-c100, mean ± SEM = −0.2 ± 0.4 cm, n = 571 neurons, p = 0.69, Wilcoxon signed-rank test).

**Figure S6:**
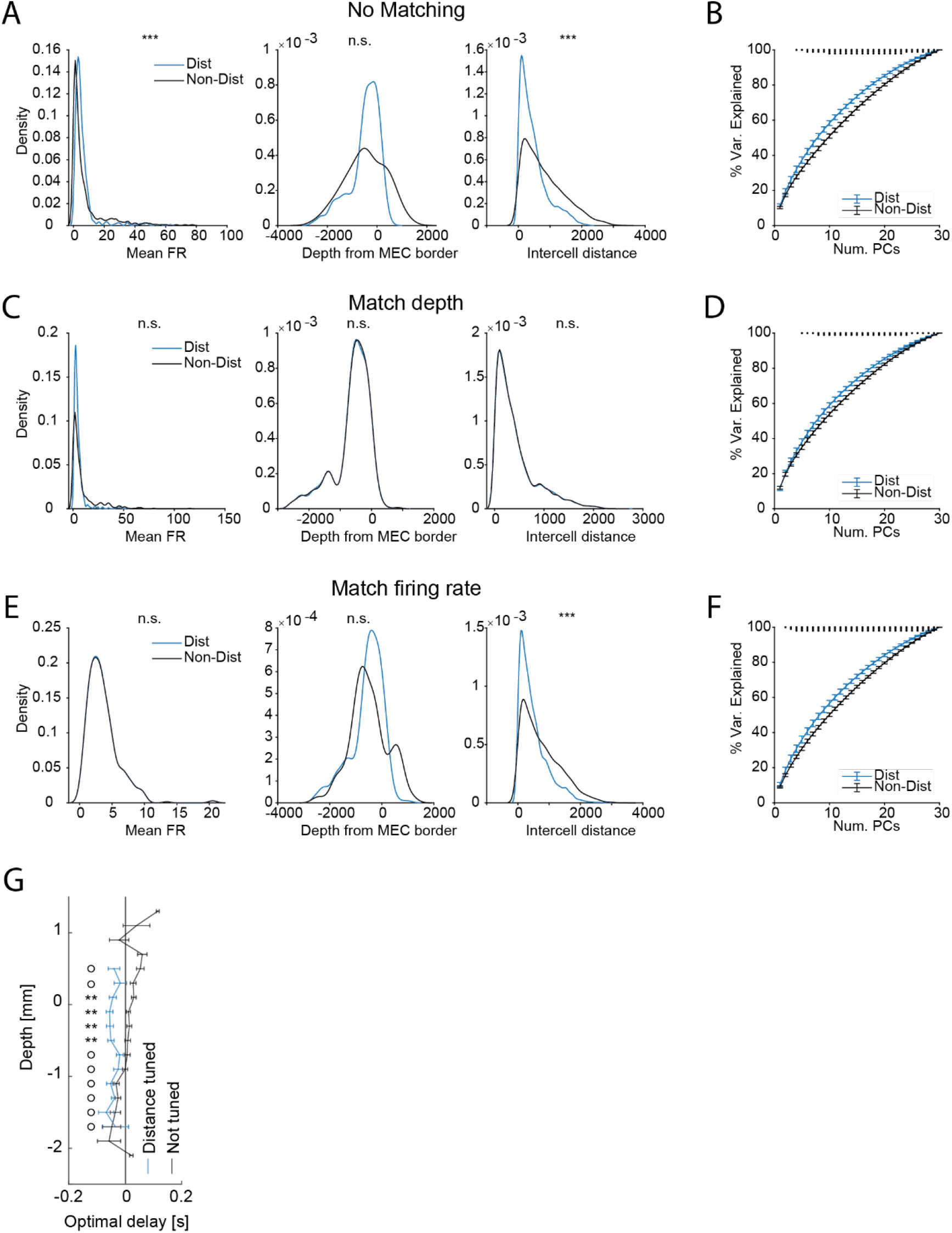
The difference in dimensionality of the activity of distance and non-distance cells is not driven by differences in depth within MEC, intercell distance, or mean firing rate. **A)** Distance neurons had different distributions of mean firing rate (left), depth within MEC (middle), and intercell distance (right) compared to non-distance neurons (mean ± SEM, mean firing rate, Hz: distance neurons = 5.6 ± 0.3, non-distance neurons = 8.1 ± 0.6, N = 450 neurons in each group, p = 2.9e-4; depth from MEC border, μm: distance neurons = −475 ± 29, non-distance neurons = −464 ± 36, N = 450 neurons in each group, p = 0.30; intercell distance, μm: distance neurons = 437 ±, non-distance neurons = 764 ± 7, N = 6525 neuron pairs in each group, p < 1e-236; Mann-Whitney U tests). These data come from 15 dark sessions in 7 mice in which at least 30 distance and 30 non-distance neurons were recorded. Thirty neurons of each type were randomly chosen for each session, resulting in a total of 30 × 15 = 450 neurons in each group, and a total of 6525 neuron pairs in each group. Neuron depths are defined as the location of peak waveform amplitude on the probe, relative to the border of MEC identified in histology. Stars above each plot show significance level of Mann-Whitney U tests (*** p < 0.001, * p < 0.05, n.s. not significant). **B)** Reproduction of Figure 6P, to compare with subsequent analyses in this figure in which neurons were matched for depth, intercell distance, and mean firing rate. Dots above each number of principal components indicate the significance of the difference in cumulative variance explained between distance and non-distance neurons (three dots, p < 0.001; two, p < 0.01; one, p < 0.05; signed rank tests; no correction for multiple comparisons). **C)** Same as (A), but non-distance neurons were chosen to match distance neurons’ distribution of depth within MEC. Specifically, for each distance neuron, we chose the closest non-distance neuron that was at least 25 μm away. As expected, matching depth on the probe also matched the distribution of intercell distances (right). Matching depth also eliminated the difference in mean firing rate (left), though the distribution of mean firing rates for non-distance neurons still had higher variance due to a longer tail of high firing rate neurons (F test p < 1e-40). **D)** Same as (B) and Figure 6P, but with non-distance neurons chosen to match depth as in (C). Even with matched depth, distance neurons had higher cumulative variance explained by small numbers of principal components. **E)** Same as (C), but with non-distance neurons to match the mean firing rate of distance neurons. Specifically, for each distance neuron, we chose the non-distance neuron with closest mean firing rate. This procedure matched distributions of mean firing rate between distance and non-distance neurons (left), but differences in the distribution of depth within MEC and intercell distance remained intact (middle, right). **F)** Same as (B) and Figure 6P, but with non-distance neurons chosen to match mean firing rate as in (E). Matching the distribution of mean firing rate between distance and non-distance neurons did not impact the result that distance neurons had higher cumulative variance explained by small numbers of principal components. **G)** Related to Figure 6N,O. Optimal temporal delay for distance tuned (blue) and non-distance tuned units (gray), averaged across sessions. For each session, neurons were assigned to overlapping depth bins (step size: 0.2 mm, bin width: 0.4 mm). For each bin, we compared the difference in optimal temporal delay factor for simultaneously recorded distance tuned and non-distance tuned units (*: p≤0.01, o: p>0.05, Wilcoxon signed rank test, bins were only included if they had data from two or more recordings). N = 23 sessions from 7 mice.

**Figure S7:**
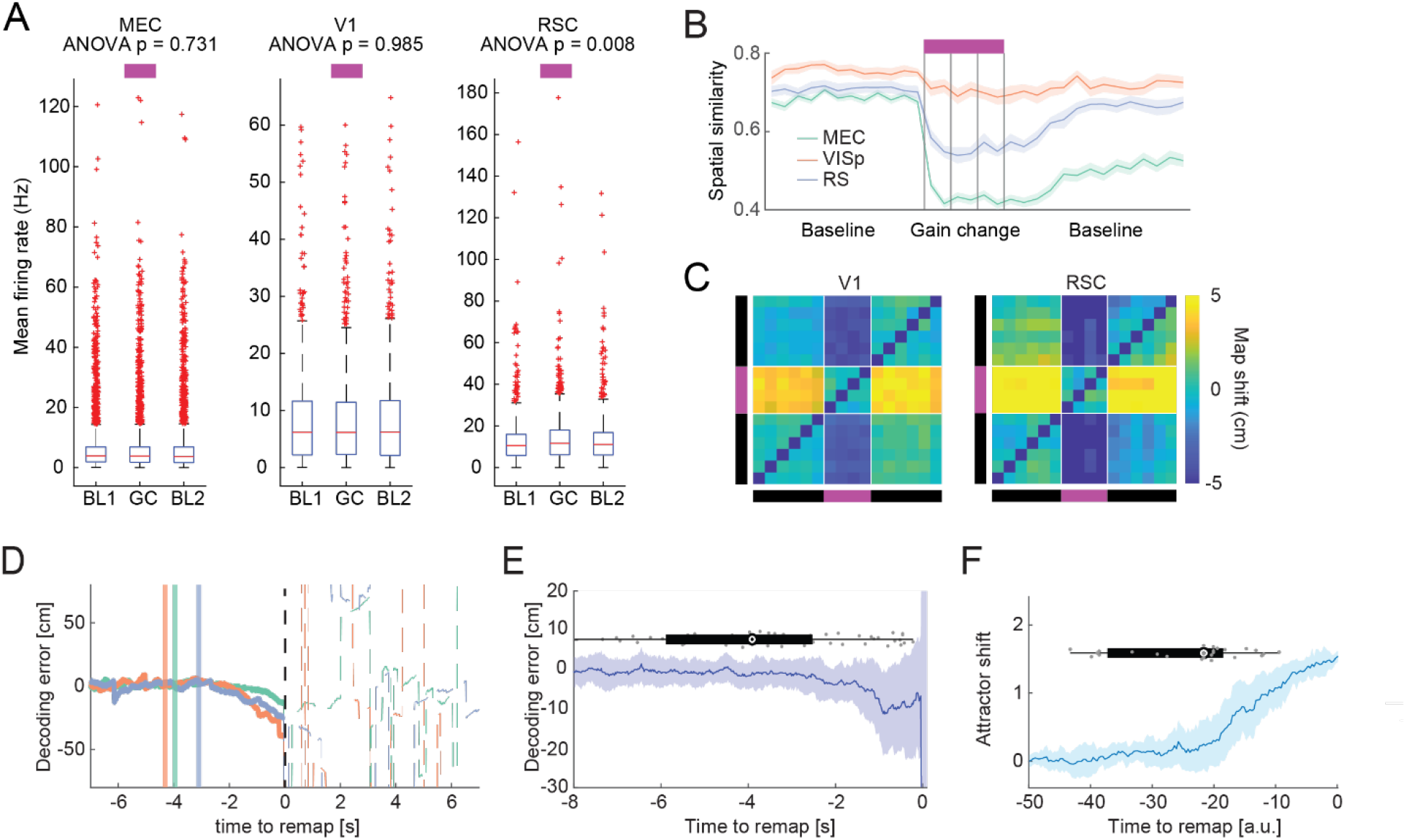
Additional characterization of responses to the gain=0.5 condition and comparison with attractor model predictions. **A)** Average firing rate increased during gain=0.5 blocks in RSC, but not in other brain regions. Only spatial cells (stability score ≥ 0.5 in baseline trials) were included in this analysis, but results were similar for all neurons (data not shown). Boxes indicate 25^th^ and 75^th^ percentiles, red line indicates median, whiskers indicate range of data points not determined to be outliers, red dots indicate outliers. **B)** Spatial similarity to baseline spatial map for blocks of 200 cm (one half of the VR track) for 6 baseline trials, 4 gain change trials (gain=0.5, trial boundaries marked by vertical lines), and 6 baseline-post trials (two data points per trial, see Methods). Lines (shading) indicate mean (SEM) across blocks. MEC: 96 blocks from 18 mice, V1: 33 blocks from 6 mice, RSC: 27 blocks from 8 mice. **C)** Average trial-trial shift matrices for V1, and RSC for gain=0.5 block. Shifts were computed as the location of the peak spatial cross-correlation for each cell, then averaged over cells within gain change blocks. White lines indicate transition between baseline and gain change trials. Black and purple bars indicate baseline and gain change trials, respectively. **D)** Decoding error as a function of time relative to remap for three examples blocks, around onset of gain=0.5. Decoder trained on firing rates of simultaneously recorded MEC neurons (see Methods). Remap was defined as time bin when |error| exceeded 50 cm. Positive/negative errors indicate a decoded position that was before/after current VR position. Solid vertical lines indicate onset of gain change. In all three examples, decoding error increases before remapping. The decoding error is consistent with map shifts observed during gain change trials. **E)** Median classification error aligned to remap for all blocks. Shading indicates median absolute deviation. Gray dots and box plot indicate distribution of gain change onsets (4 data points outside axis limits). **F)** Similar to (E), but for data simulated based on coupled attractor model. The sign of the attractor shift is opposite to the sign of decoder error.

## Methods

### Experiments

#### Subjects

All techniques were approved by the Institutional Animal Care and Use Committee at Stanford University School of Medicine. Recordings were made from 32 female C57BL/6 mice aged 12 weeks to 6 months at the time of surgery (18 −26 grams). Mice were housed singly or with littermates in transparent cages on a 12-hour light-dark cycle and experiments were performed during the light phase.

#### Surgery

Anesthesia was induced with isoflurane (4%; maintained at 1.75%) followed by injection of buprenorphrine (0.1 mg/kg). Fiducial marks were made on the skull ± 3.3 mm lateral from the midline and approximately 4 mm posterior from Bregma, and ± 2.5 mm lateral from the midline and approximately 3 mm posterior from Bregma for targeting V1 and RSC. A ground screw was affixed to the skull approximately 2.5 mm anterior and 1.5 mm lateral from Bregma on the left side. A stainless steel headbar was attached to the skull using Metabond and the rest of the exposed skull was covered with Metabond. Transparent Metabond was used to allow visualization of the fiducial marks, which would later guide craniotomies and probe placement. After training, which typically lasted 2-3 weeks, mice were again anesthetized with isoflurane and the Metabond and skull were shaved down with a dental drill, posterior to the fiducial mark on both sides. For MEC recordings, bilateral craniotomies, roughly 500 μm in diameter, were made posterior to the fiducial mark, exposing the transverse sinus. Plastic rings cut from pipette tips (~4 mm diameter) were affixed to the skull around each craniotomy using Metabond. These rings served as wells to hold saline and a layer of silicone oil during recording, which prevented the craniotomy from drying out. Craniotomies were covered with KwikCast and the mouse recovered overnight before the first recording session. A maximum of three recording sessions were performed per hemisphere, each on a different day, for a maximum of six recording sessions on six days total for all mice but one (in which seven recordings sessions were performed across seven days). Durotomies were performed before recording from a craniotomy for the first time, either the same day as the craniotomy (side 1, typically the right side) or three days later (side 2, typically the left side). For recordings targeting V1 and RSC, we followed a similar strategy, except that only one craniotomy, combined with a durotomy, was performed at a time. We then performed 2 recordings per craniotomy before targeting a new location with another brain region. As for MEC recordings, a maximum of 6 recordings per mouse was performed.

#### Virtual reality setup

The VR setup was the same as in (Campbell et al., 2018), except that the VR scene was displayed on three 24-inch monitors surrounding the mouse instead of a 14-inch hemispherical projection screen, and the mouse ran on a 6-inch diameter foam cylinder instead of a 20-cm diameter Styrofoam ball. Briefly, rotation of the treadmill was recorded using a rotary encoder (Yumo, 1024P/R), and data was processed and transmitted to the virtual reality software via a microcontroller (Arduino UNO) using custom software. The virtual reality environment was created and displayed using custom software in Unity 3D (Unity Technologies). The training environment consisted of a floor with checkerboard texture and evenly spaced, identical visual landmarks on both sides. The test environment had the same floor as the training environment, and in addition to the reward tower (which remained the same), contained 4 additional visual landmarks (towers). These five visual landmarks (towers) were identical on the left and right sides and were evenly spaced at 80 cm intervals, covering a total of 400 cm. Upon reaching the tower at 400 cm, mice were ‘teleported’ to the beginning of the track for the next trial without any visual discontinuity, giving the impression of running along an infinite hallway. Water rewards were delivered via Tygon tubing attached to a metal lick spout mounted in front of the mouse. Delivery was triggered via a solenoid valve, which produced an audible clicking sound upon reward delivery. Licking was detected with a custom-built infrared light barrier. The silicon probes were mounted either on a manual manipulator on a custom rotatable mount (World Precision Instruments) or a motorized micromanipulator (UMP Micromanipulator, Sensapex). Probe holders were placed behind the mouse to minimize visual interference.

#### Behavioral training

After headbar implantation, mice recovered for three days and were given Baytril (10 mg/kg) and Rimadyl (5 mg/kg) daily. After three days, they were taken off Baytril and Rimadyl and put on water deprivation. They received 0.8 mL of water each day and their weights were monitored to ensure that they remained above 85% of baseline. Training progressed in three stages. In stage one, they were head-fixed on the VR rig and trained to receive water from the lickspout. Water delivery was associated with an audible click of the solenoid. Mice quickly learned this association and began licking upon hearing the click. After stage one, which typically lasted two to four days, mice progressed to stage two, in which they ran on a training track to receive water rewards (see *Virtual Reality Setup* for description of training track). To encourage mice to run, water rewards (~2 μL) were automatically delivered whenever they passed a tower. The tower spacing started at 40 cm and was increased daily to encourage running, up to a maximum of 200 cm. The reward tower on the habituation track was visually identical to the reward tower on the track used for recording. Once mice ran consistently on the habituation track (average running speed > 10 cm/s), they progressed to stage three, in which they ran on the same track that would eventually be used for recording. They ran 50 – 200 trials per day for 4 – 20 days (median 14 days) for a total of 800 – 2500 trials (median 2000 trials) before the first recording session. During this final phase of training, mice developed stereotyped running and licking patterns in which they slowed down and licked prior to the reward tower. Some training sessions were performed on a training rig, but mice were always trained on the recording rig for several days prior to the first recording day to familiarize them with the setup.

#### Electrophysiological recording

Extracellular recordings were made by acutely inserting silicon probes into a craniotomy above the target brain region while the animal was head-fixed in the VR setup. One insertion was performed per day per mouse, after which the craniotomy was covered with Kwik Cast and returned to on subsequent days for additional recordings. Typically, we performed 3 insertions on each side for MEC recordings, and 2 insertions per craniotomy for recordings targeting V1 and RSC (all in similar sites within the same craniotomy) for a total of 6 recordings per mouse over 6 days.

Most recordings were made using single Neuropixels probes (Jun et al., 2017; Lopez et al., 2016). For MEC recordings, 17 recordings from 7 mice were made with Phase 3A Option 4 probes (276 recording channels), and 62 recordings from 11 mice were made with Neuropixels 1.0 probes (384 recording channels, www.neuropixels.org). For recordings targeting V1 (24 sessions, 9 mice) and RSC (18 sessions, 9 mice), all recordings were made with Neuropixels 1.0 probes. Raw voltage traces were filtered, amplified, multiplexed, and digitized on-probe and recorded using SpikeGLX (https://billkarsh.github.io/SpikeGLX). Voltage traces were filtered between 300 Hz and 10 kHz and sampled at 30 kHz with gain = 500 (AP band) or filtered between 0.5 Hz and 1 kHz and sampled at 2.5 kHz with gain = 250 (LFP band). All recordings were made using the contiguous set of recording channels closest to the tip of the probe (Bank 0). The probe’s ground and reference pads were shorted together and soldered to a gold pin, which was then connected to a gold pin that had been soldered to a skull screw on the animal. For Phase 3A Option 4 probes, recordings were made in external reference mode, using the skull screw as external reference. For Neuropixels 1.0 probes, recordings were made in internal reference mode, with the probe tip set as reference, but the ground and reference pads were still connected to the skull screw in order to ground the animal and prevent electrostatic damage to the probe.

In some recordings (11 recordings from 3 mice, 597 units total), we used 64-channel silicon probes from Cambridge Neurotech (model H3, www.cambridgeneurotech.com). Data were recorded using a Neuralynx Digital Lynx SX data acquisition system.

Prior to insertion, probes were first dipped in one of three different colors of dye (DiI, DiD, and DiO). Different colors were used to allow up to three different penetrations in the same craniotomy to be distinguished in histology. The mouse was head-fixed on the VR rig and the KwikCast was removed from above the craniotomy. For MEC recordings the probe was mounted on the rig at a slight angle to maximize alignment between the probe and MEC (approx. 10 degrees). The probe was then lowered down to the level of the skull, aligned mediolaterally with the fiducial mark, and inserted as close to the transverse sinus as possible. The well was filled with saline (0.9% NaCl) and covered with a layer of silicone oil to prevent drying. The probe was inserted slowly (~10 um/sec) until there was no longer any activity at the tip or until the probe started to bend. The probe was retracted 100-200 μm from its deepest penetration point and left to settle for 30 minutes before starting the recording. For recordings targeting V1 and RSC, we followed a similar strategy.

Recordings lasted between 40 and 150 minutes, during which the mouse ran one or more VR sessions under various gain, contrast, and cue removal conditions. The VR emitted a TTL pulse every frame from an Arduino UNO, and these pulses were recorded in SpikeGLX using an auxiliary National Instruments data acquisition card (NI PXIe-6341 with NI BNC-2110) to synchronize VR traces with neurophysiological data.

After each recording session, the probe was rinsed with deionized water, soaked in deionized water for at least 15 minutes, soaked in an enzymatic detergent (2% Tergazyme in deionized water) for at least 60 minutes, and rinsed and soaked again in deionized water for at least 15 minutes.

#### Dark and passive playback recordings

There were two separate experiments in which mouse ran in the dark. In the first experiment, mice ran in the dark with rewards delivered every 400 cm (baseline trials with monitors turned off; Figure S1F,G). These dark sessions occurred in between two VR sessions with the monitors on. In the second experiment, mice ran freely in the dark for ~30 minutes and no rewards were delivered (Figure 6). These dark sessions occurred after mice ran VR sessions with gain or contrast manipulations (~200 trials).

In a separate set of experiments, mice passively sat on the running wheel while visual stimuli from a previous VR session were played back (Figure S1D,E). The running wheel was braked during passive playback. Mice used in these experiments were habituated to the braked condition during training. Passive playback sessions occurred in between two 50-trial VR sessions. The first 50-trial VR session was used to generate the playback stimuli.

#### Histology

After the last recording session, the mouse was euthanized with an injection of pentobarbital (0.2 ml i.p. of 100mg/ml beuthanasia) and perfused transcardially with 1X phosphate-buffered saline (PBS) followed by 4% paraformaldehyde (PFA). Brains were extracted and stored in 4% PFA for at least 24 hours after which they were transferred to 30% sucrose and stored for at least 48 hours. Finally, brains were frozen and cut into 65 μm sections (sagittal sections for MEC recordings, coronal sections for V1 and RSC recordings) with a cryostat, stained with DAPI, and imaged with a widefield fluorescence microscope (Axio Imager 2, Zeiss). To localize MEC recordings we identified the sections where the probe entered MEC and where the tip of the probe was localized. From these two points, we determined the section of the probe within MEC and only used units recorded within this span for further analysis. To reconstruct the probe tracts for V1 and RSC recordings, we used software written in MATLAB (https://doi.org/10.1101/447995) (Shamash et al., 2018). This allowed us to first register the coronal sections to a standard atlas and then reconstruct the probe tract within the standard atlas framework.

### Data analysis

All analyses were performed in MATLAB using custom scripts.

#### Spike sorting and synchronization

Spike sorting was performed offline using Kilosort2 (https://github.com/MouseLand/Kilosort2) (Pachitariu et al., 2016). Data from the Phase 3A Option 4 Neuropixels probes and the Cambridge Neurotech H3 probes, which were gathered prior to the release of Kilosort2, were sorted with Kilosort1 (https://github.com/cortex-lab/KiloSort). Clusters were manually inspected and curated in Phy (https://github.com/cortex-lab/phy). Spike times and cluster identities were extracted from the output of KiloSort2 and Phy using code from the Spikes repository (https://github.com/cortex-lab/spikes). VR data (VR position, lick times, and contrast and gain values for each trial) were synchronized to spiking data by substituting the time of each VR frame with the time of the corresponding detected TTL pulse. Synchronization was checked by comparing the difference between subsequent VR frame times with the difference between subsequent TTL pulse times and confirming that these were highly correlated (*ρ* > 0.95). For data acquired with Neuropixel 1.0 probes, we additionally corrected any drift between the IMEC PXIe acquisition module and the auxiliary acquisition module (NI PXIe-6341) by aligning the sync waveforms generated by the IMEC PXIe. This was necessary as the two modules tended to drift apart by about 5ms/hr. Because the frame rate of the VR was not constant but instead fluctuated slightly around 60 Hz, we used linear interpolation to resample VR data to a constant 50 Hz for ease of subsequent analysis. Finally, spike times, cluster identities, and synchronized VR data were saved together in a single file for analysis in MATLAB (one file per VR session).

#### Spatial firing rate maps

To compute spatial firing rate maps, we binned the position into 2cm bins, computed spike counts for each position bin, divided the counts by the occupancy of each spatial bin and smoothed the resulting spatial firing rate (Gaussian kernel, standard deviation = 4 cm). To calculate average firing rate as a function of position (Figure 1J), we first averaged the spatial firing rate maps across trials for a neuron and across spatially stable neurons for a recording session. We then finally averaged these per-session firing rates, excluding blocks with fewer than 5 stable neurons (see below), for each brain region.

#### Running speed and lick rate

To calculate the running speed of the mouse, we calculated the difference in VR position between consecutive VR frames. The resulting running speed trace was smoothed with a gaussian kernel (sigma = 0.2 s). To calculate running speed as a function of position, we averaged the smoothed running speed for each position bin. We measured licking behavior with an infrared light barrier. Thus, licking resulted in drops of photodiode voltage output. Photodiode voltage was monitored by an Arduino UNO. The VR computer queried the voltage on each frame and individual licks were defined as the voltage dropping below a predefined threshold.

#### Spatial stability and similarity matrix

To compute the spatial stability measure and the spatial similarity matrix for a neuron, we first computed trial by trial spatial firing rate maps as described above, subtracted the mean firing rate from each trial, and then computed the normalized cross correlation with a maximal lag of 30 cm for each pair of trials. The similarity for a pair of trials was defined as the maximum value of this cross correlation. To compute a spatial stability score for a given set of trials for each cell, we averaged all pairwise cross correlations. We defined a neuron as stable if the resulting average was ≥ 0.5. To calculate a similarity matrix for a block of trials in a recording session, we averaged individual similarity matrices of each neuron that fulfilled the stability criteria for a set of trials (usually the 6 trials preceding a gain change). Note that in this way, neurons did not have to fulfill the stability criteria across the whole session. Blocks with fewer than 5 cells that fulfilled the stability criteria were excluded.

To quantify trial-by-trial similarity as a function of inter trial distance (Figure S2H), we calculated pairwise stability as described above for all pairs of co-recorded units for a block of 16 baseline trials. We included only units with average spatial stability ≥ 0.5 across pairs of neighboring trials.

To calculate spatial similarity of baseline maps (Figure S2I), we first calculated the spatial firing rate maps for each block by averaging across 6 trials before the onset of the gain change. For each unit, we then calculated the spatial similarity as described above between these two blocks and averaged across all co-recorded units of a block. We only included units that were stable in both baseline blocks, i.e. the average spatial similarity was ≥ 0.5 in both blocks of baseline trials.

To calculate spatial similarity for blocks of 200 cm (Figure S7B), we first calculated the baseline firing rate maps for each block by averaging across 6 trials before onset of the gain change. For each unit, we then calculated the spatial similarity for each section of 200 cm by calculating the cross correlation between the mean subtracted spatial firing rate of the current section, and the corresponding baseline section. Note that, for gain change trials and post-baseline trials, the baseline map was calculated by averaging across all 6 pre-baseline trials. For the similarity to baseline for pre-baseline trials, the baseline spatial firing rate map was obtained by averaging across the remaining 5 pre-baseline trials.

#### Map shift

To calculate the shift between spatial firing rate maps, we computed spatial similarity matrix as described above. We estimated the map shift between pairs of trials as the location of the maximum cross correlation. As for the similarity matrices, we averaged these shift matrices across all stable neurons of a block to estimate the population trial-by-trial map shifts. To estimate the average shift between baseline trials and gain change trials, we averaged the population shift across all pairs of baseline trials and gain change trials (indicated by red rectangle in Figure 3C).

#### Position decoding

To decode position from the firing rates of simultaneously recorded neurons (Figure 1K, Figure S3F, Figure 7G, Figure S7E) we first chose a set of encoding (training) trials. Cells were included for analysis if they had a stability score ≥ 0.5 in encoding trials (see above for definition of stability score). For these neurons, we calculated the z-scored firing rate over time (time bin: 0.02 s, Gaussian smoothing sigma: 0.2 s) and used the z-scored firing rate to compute spatial tuning curves for encoding trials by averaging the population activity within each position bin (2 cm bins). This defined an average N-dimensional trial trajectory during encoding trials, where N is the number of neurons. For each time point during decoding (test) trials, we determined the closest point on the average encoding trajectory by minimizing the Euclidean distance. The decoded position then corresponded to the VR position for this point on the average trajectory. In both encoding (training) and decoding (test) data, we only considered times when the mouse was running (cutoff: 2 cm/s).

We used this approach to study the time course of MEC remapping in response to gain = 0.5 (Figure S7D, E, F). We inferred the location and time of remapping by finding the first instance when the absolute decoding error became larger than 50 cm, preceded by a period of at least 3 seconds with absolute error smaller than 50 cm.

We also used this approach to estimate the dependence of decoded position on running speed (Figure S3F). For blocks of 10 baseline trials with at least 5 stable cells (stability score ≥ 0.5), we decoded position in each trial by treating the other 9 trials as the encoding trials. We computed decoding error as the difference between decoded position and actual position, respecting the circularity of the track. For example, if decoded position were 399 cm and true position were 1 cm, the error would be −2 cm, not 398 cm. We removed times when this difference exceeded 50 cm. We concatenated decoder error, true position, and running speed traces across trials, and used multiple linear regression to estimate the influence of running speed and position bin (bin size = 20 cm) on decoder error. For each session, we averaged coefficients over all 10 trial blocks with at least 5 stable cells (Figure S3F). The sign of the coefficient was flipped in Figure S3F to match the signs of retrospective/prospective coding conventions in Figure 4.

To determine the decoding error as a function of number of included neurons in baseline trials (Figure 1K), we followed a similar decoding procedure. First, we calculated spatial firing rate maps for trials 5-20 for each neuron in each recording session. Note that these were always baseline trials. We excluded the first 4 trials because we noted that spatial firing rates tended to be more variable at the beginning of a recording. For further analysis, we only included neurons with a stability score ≥ 0.5 across all 16 trials.

To estimate decoding accuracy for N included neurons, we drew a random sample of spatial maps of N neurons. We then averaged the spatial firing rates across 15 training trials for each neuron (leaving out one trial for test). This gave us the training firing rate trajectory in N dimensions. To decode the position from the test trial, we found the closest point on the training trajectory for each position bin by minimizing the Euclidean distance, as above. In this way, we found the decoded position for each position bin of the test trial. We repeated this procedure for all 16 trials. This gave us an estimated position for each position bin in each of the 16 trials. We computed the decoding error by taking the average of the absolute difference between actual position and decoded position, across 16 trials, respecting the circularity of the track as above. To estimate the distribution of the absolute error, we also calculated the histogram of the absolute error. We repeated this approach for each set of N neurons by drawing random sets of N neurons 100 times.

#### Clustering of similarity matrices

To cluster the similarity matrices (Figure 2F), we first vectorized the upper half of each similarity matrix and stacked the vectorized trial blocks into a matrix, resulting in a N x M matrix, where N is the total number of trial blocks across all three brain regions, and M is the number of trial pairs. We then calculated the principle components of this matrix and used the first three principal components to cluster the blocks into 3 clusters with k-means clustering with 3 clusters. We used 3 clusters because this corresponded to the elbow of the plot of total distance to centroids versus number of clusters (Figure S2G). To estimate the remapping behavior in baseline trials in MEC, we calculated spatial similarity matrices of blocks of 16 baseline trials, including only neurons that had a spatial stability score ≥ 0.5 across the first 6 trials. We then vectorized the upper half of each similarity matrix as described above and subtracted from each vector its mean to center the data, and projected onto the first three principal components of the gain change block similarity matrix to get a projection score for each baseline-only block. We assigned each baseline-only block to the closest cluster, by calculating the minimum distance between the centroid of each cluster and the projection scores.

#### Estimating coordination of remapping

To calculate within block correlation between the spatial similarity map of individual units and the remainder of the co-recorded population (Figure 2), we calculated the correlation between the spatial similarity map of each unit and the population similarity map. The population similarity map was obtained by averaging across the remaining group of simultaneously recorded spatial units. This was repeated for each unit and then averaged across all co-recorded spatial units for a block to obtain an average within-block correlation of the individual spatial similarity maps to the population similarity map. To calculate across-block correlation, we correlated the spatial similarity map of each unit for a block, with the population maps of all other ‘remap’ blocks, and averaged across all co-recorded spatial units to obtain the average across-block correlation.

#### Estimating temporal delay factors

To visualize the effect of running speed on spike location (Figure 4C), we calculated the average spatial firing rate maps for each neuron as described above for blocks of 16 baseline trials. Gain change trials were not included in this average. We identified the VR position with maximum average firing rate for each cell (indicated by a red line in Figure 4C). Finally, we created raster plots for each neuron in a region surrounding the maximum firing rate location (−50 cm to + 40 cm) sorting trials by the average running speed (average over 10 cm preceding location of maximum firing rate).

To estimate the temporal delay for a neuron, we employed a grid search over a range of values *d* (−300 ms to 300 ms, in 10 ms increments). We calculated time shifted spatial firing rate maps for each trial as described above, but for 1cm spatial bins and a smoothing kernel with sigma=3.5 cm, and shifting the time of each spike by *d*. To estimate the trial by trial alignment of firing rate maps, we computed trial-by-trial correlations of the time-shifted spatial firing rate maps for each pair of trials. To minimize boundary effects, we only used positions between 10 cm and 390 cm. We estimated the temporal delay *d* for a neuron for a block of trials as the value *d* that resulted in the highest average correlation across trials (e.g. Figure 4D). As for map shifts, we ignored *d* for a block of trials if the maximum occurred at either limit. To reduce overfitting and minimize the effects of remapping and drifts in spatial firing, we repeated this procedure for blocks of continuous baseline trials (blocks were non-overlapping). For each neuron, the final estimate of temporal delay was the average across multiple blocks (8 – 18 blocks, median: 11, depending on trial structure of the recording session). Blocks of trials were ignored if the spatial stability of the neuron was smaller than 0.5. Values were used for further calculations if at least 3 blocks fulfilled the range criterium (*d* in (−300, 300)) and stability criterium (MEC: 26% of neurons, V1: 38% of neurons, RSC: 45% of neurons). The same approach was used to estimate temporal delay at different contrast levels (Figure S5G).

#### Map shift after temporal delay correction

For each block of the ‘stable’ cluster, we shifted the spike times of each neuron by its estimated temporal delay, and calculated similarity matrices, shift matrices and map shifts as described above. We then calculated the difference in map shift between uncorrected and corrected pairs of blocks (Figure 4F).

#### Spatial firing rates in low contrast

In a subset of mice, we reduced the contrast of the visual landmarks to test the influence of visibility of the landmarks on neural activity. We started with a block of 24 trials of high contrast. Two sets of 10 trials with baseline gain were interleaved with gain = 0.8. This was followed by a block of 24 trials with low contrast, with same gain structure as for the high contrast block. This allowed us to test the influence of visual contrast on neural firing both during baseline gain as well as gain = 0.8. We calculated the shift of spatial maps during gain = 0.8 as described above, including only trials with spatial similarity ≥ 0.5 to the baseline map, and only considering neurons with spatial stability ≥ 0.5 in the 6 trials preceding the gain change both during high contrast and low contrast (Figure 5D). To visualize the effect of contrast on the location of peak firing rate (Figure 5H), we calculated the average spatial firing rate maps for each neuron as described above for blocks of 6 trials preceding gain change in high contrast. We then determined the location of peak firing and averaged spatial firing rates during gain change trials around the location of peak firing rate (+/−30 cm), normalizing to the maximum firing rate within each trial. We repeated this for gain change during low contrast. We then averaged the peak firing rate curves for high and low contrast for all neurons with spatial stability ≥ 0.5 in the 6 trials preceding the gain change both during high contrast and low contrast.

#### Spatial autocorrelation and distance tuning

To compute the spatial autocorrelation of firing rate traces during dark sessions (Figure 6), we first computed the firing rate as a function of distance, similar to the spatial firing rate but replacing position by distance run since start. We then computed the autocorrelation of this firing rate trace, including lags up to 800 cm. To identify the preferred distance of a neuron, we used MATLAB’s *findpeaks* function to detect the most prominent peak in the spatial autocorrelation. In addition, we computed a shuffled spatial autocorrelation by shifting spike times relative to elapsed distance by random offsets (n=300 shuffles, offset drawn from a uniform random distribution with the interval (20 seconds, max_t), where max_t was the duration of the recording). For each shuffle, we computed the maximum autocorrelation peak up to a maximum lag of 800 cm. We determined that a neuron was distance-tuned if the prominence of the peak was greater or equal 0.1, and the height of the autocorrelation at this preferred distance was greater than the 99^th^ percentile of the shuffled distribution of autocorrelation peaks. This threshold was used to compare gain change responses, temporal delay factors, and dimensionality between distance and non-distance neurons (Figure 6 J-Q).

#### Estimating dimensionality of distance and non-distance cell activity using principal components analysis

We used dimensionality reduction with PCA to compare dimensionality of distance and non-distance cell activity during running in the dark (Figure 6Q). We first chose dark sessions with at least 30 of each cell type and randomly selected 30 distance and non-distance cells for each of these sessions. We computed smoothed firing rate traces for each neuron (time bin = 20 ms, standard deviation of Gaussian filter = 200 ms), z-scored each neuron’s firing rate trace separately, and performed PCA on the resulting firing matrix (separately for distance and non-distance cells). We compared cumulative variance explained by the top N principal components between distance and non-distance cells. To control for differences in mean firing rate and anatomical location between distance and non-distance cells, we performed the same analysis but chose, for each distance neuron, the non-distance neuron with the closest mean firing rate or anatomical location (requiring a minimum distance of 25 um between cells to avoid cross-contaminated clusters) (Figure S6).

### Modeling

#### Kalman filter

The Kalman filter is a method for tracking variables of interest by balancing external inputs with updates from an internal model describing how they evolve over time. The Kalman filter assumes that the variables are normally distributed and the processes updating the internal estimate are linear. The assumption that variables are Gaussian is convenient because it means we only have to track their means and variances to fully describe them, and the assumption that the internal process is linear makes it easy to model. Importantly, when these assumptions are met, the Kalman filter is the optimal Bayesian solution to this problem.

We consider the evolution of a position estimate, *x_A_*, which is updated by both internal path integration and external input from landmarks located at *x_L_* according to a continuous Kalman filter. In the Kalman filter formulation, the internal estimate *x_A_* and external inputs *x_L_* both have associated uncertainties, which we call *σ_A_* and *σ_L_*. Going forward, it is mathematically convenient to work with the *difference* between *x_A_* and *x_L_*, which we call Δ*x* = *x_A_* − *x_L_*. We now consider how Δ*x* evolves in time according to a Kalman filter.

In the Kalman interpretation, the relative strengths of landmark and self-motion inputs are determined by the different levels of uncertainty attached to them. In the real world, these uncertainties correspond to the magnitudes of random errors in landmark perception and path integration. Here, for simplicity, we ignore these random errors and only consider the levels of uncertainty. When random noise is incorporated into the model, the distribution of position self-estimates will be a Gaussian distribution centered around the results derived here.

To build up to a continuous Kalman model, we first consider a series of closely spaced, weak landmark input events, separated by time Δ*t*. At each landmark event, the landmark input tells the system it is at Δ*x* = 0 with some uncertainty *σ_L_*. Because the landmark events are frequent and weak, we have *σ_L_* ≫ *σ_A_*. The position estimate Δ*x* becomes a weighted average of the original position estimate (Δ*x*) and the position estimate given by new landmark information (Δ*x* = 0) where the relative weighting is given by the uncertainties associated with each signal:

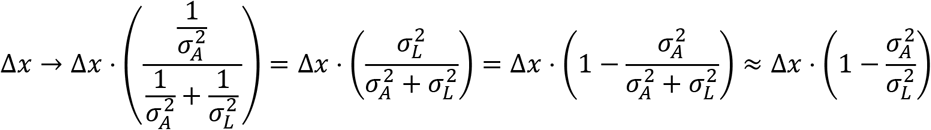

And the uncertainty *σ_A_* of the position estimate after the landmark event will decrease in light of the increased information as follows:

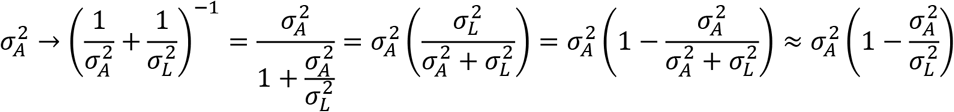

In both cases, the last simplification was made using the fact that *σ_L_* ≫ *σ_A_*.

In between the landmark events, the animal performs linear path integration:

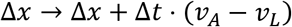

where *v_L_* is the velocity of the landmarks and *v_A_* is the animal’s internal velocity estimate. For simplicity, we are not modeling noise in the velocity estimate. When there is no gain change, *v_A_* − *v_L_* = 0. During a gain change *v_A_* differs from *v_L_* and so this term is non-zero (Figure S4C).

We model the position uncertainty to grow linearly with time according to a diffusivity constant *α*:

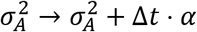

Now we can take the limit Δ*t* → 0. We can write the landmark uncertainty as 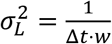 where *w* is the “Kalman landmark strength” and has units of 1/(*cm*^2^ ⋅ *time*). In this limit, the difference equations become:

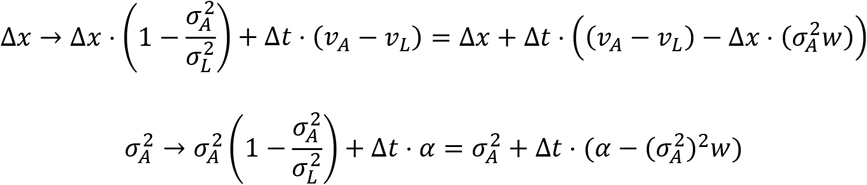

The second equation reaches an equilibrium at

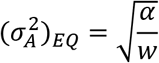

Which makes the first equation reach equilibrium at

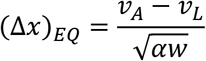

This steady state solution is exactly analogous to the linearization of the sub-critical regime of the coupled oscillator model around *D* = 0, where 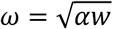.

While a Kalman filter captures the equilibrium behavior of the attractor model’s sub-critical regime, it differs from the attractor model in two key ways. First, the attractor model as it stands has no representation of attractor uncertainty. One implication of this difference is that the two models make different predictions about the impact of landmarks on the future behavior of the system after the landmarks have been encountered. To see this, note that the Kalman model only reduces to the attractor model when we assume 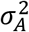 has reached an equilibrium, and to reach equilibrium, the Kalman model requires uniform values of 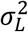. In reality, different landmarks can have different strengths. In this situation, the Kalman model and the attractor model make different predictions. If a Bayesian agent observes very strong landmarks, its certainty about its position self-estimate will go up, and it should weight subsequent landmarks less. Thus, if the animal encounters a particular strong/weak landmark during a gain change, 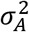 will decrease/increase, causing *future* landmarks to have less/more of an effect on the position estimate. The attractor model does not have any representation of uncertainty, so it does not predict this memory-like effect.

Second, because a Kalman filter is linear, it does not yield a super-critical regime. Instead of de-cohering after a sharp threshold, the position estimate will continue to shift indefinitely. Note that the equilibrium value of the position uncertainty 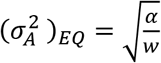 only depends on the diffusivity *α* and the Kalman landmark strength *α*. Thus, for the Kalman model, all values of gain will lead to pure shifts of the firing patterns, without any remapping, rescaling, or warping of any kind. Therefore, the Kalman model only applies to the “stable” (Cluster 1) responses (Figure 2G) and cannot capture the remapping phenomena we observed in the data (Figure 2H-J, Figure 7).

